# Higher-order thalamocortical inputs gate synaptic long-term potentiation via disinhibition

**DOI:** 10.1101/281477

**Authors:** Leena E. Williams, Anthony Holtmaat

## Abstract

Sensory experience and perceptual learning changes the receptive field properties of cortical pyramidal neurons, largely mediated by long-term potentiation (LTP) of synapses. The circuit mechanisms underlying cortical LTP remain unclear. In the mouse somatosensory cortex (S1), LTP can be elicited in layer (L) 2/3 pyramidal neurons by rhythmic whisker stimulation. We combined electrophysiology, optogenetics, and chemogenetics in thalamocortical slices to dissect the synaptic circuitry underlying this LTP. We found that projections from higher-order, posteriormedial thalamic complex (POm) to S1 are key to eliciting NMDAR-dependent LTP of intracortical synapses. Paired activation of intracortical and higher-order thalamocortical pathways increased vasoactive intestinal peptide (VIP) interneuron and decreased somatostatin (SST) interneuron activity, which was critical for inducing LTP. Our results reveal a novel circuit motif in which higher-order thalamic feedback gates plasticity of intracortical synapses in S1 via disinhibition. This motif may allow contextual feedback to shape synaptic circuits that process first-order sensory information.

## INTRODUCTION

Sensory experience and perceptual learning can remodel neocortical synaptic circuits throughout life (Feldman, 2009). The long-term potentiation and depression of synapses (LTP and LTD, respectively) constitutes a fundamental underpinning of functional cortical synaptic circuit plasticity (Bliss and Collingridge, 1993; Sjöström et al., 2008; Feldman, 2009; Froemke, 2015). However, the circuit mechanisms of cortical LTP and LTD remain unclear. In particular, the interactions of long-range feedback projections with local cortical microcircuits, and the role thereof in local cortical plasticity have been poorly investigated.

The mouse somatosensory cortex (S1) serves as an important model for LTP and LTD, largely owing to the one-to-one anatomical relationship between individual sensory organs (whiskers) and the cortical columns (Feldman, 2009). Hence, it is relatively easy to perform targeted recordings, as well as to selectively enhance or decrease sensory input. First-order, whisker sensory information passes to S1 through the ventroposterior medial (VPM) thalamus which projects onto layer (L) 4 and L5b, representing the lemniscal pathway (**Figures 1A,B**) (Feldmeyer, 2012). L4 and L5b neurons in turn synapse, among others, onto L2/3 pyramidal neurons (Lefort et al., 2009; Petreanu et al., 2009; Feldmeyer, 2012). Higher-order thalamocortical feedback from the posteromedial thalamic complex (POm) joins ascending sensory input to S1 and projects onto L2/3 and L5a neurons, representing the paralemniscal pathway (Bureau et al., 2006; Petreanu et al., 2009; Feldmeyer, 2012; Jouhanneau et al., 2014). Therefore, both lemniscal inputs (via L4) and paralemniscal inputs (via direct POm projections) arrive at L2/3 pyramidal neurons. L2/3 pyramidal neurons are inhibited by a variety of interneurons. In particular, their distal dendrites are strongly inhibited by somatostatin (SST)-expressing interneurons (Wang et al., 2004; Gentet et al., 2012), which, in turn, are inhibited by vasoactive intestinal peptide (VIP)-expressing interneurons (Pfeffer et al., 2013; Lee et al., 2013). The lemniscal (L4) and paralemniscal (POm) pathways provide direct and indirect input to both interneuron types (Wall et al., 2016; Audette et al., 2017).

**Figure 1.**
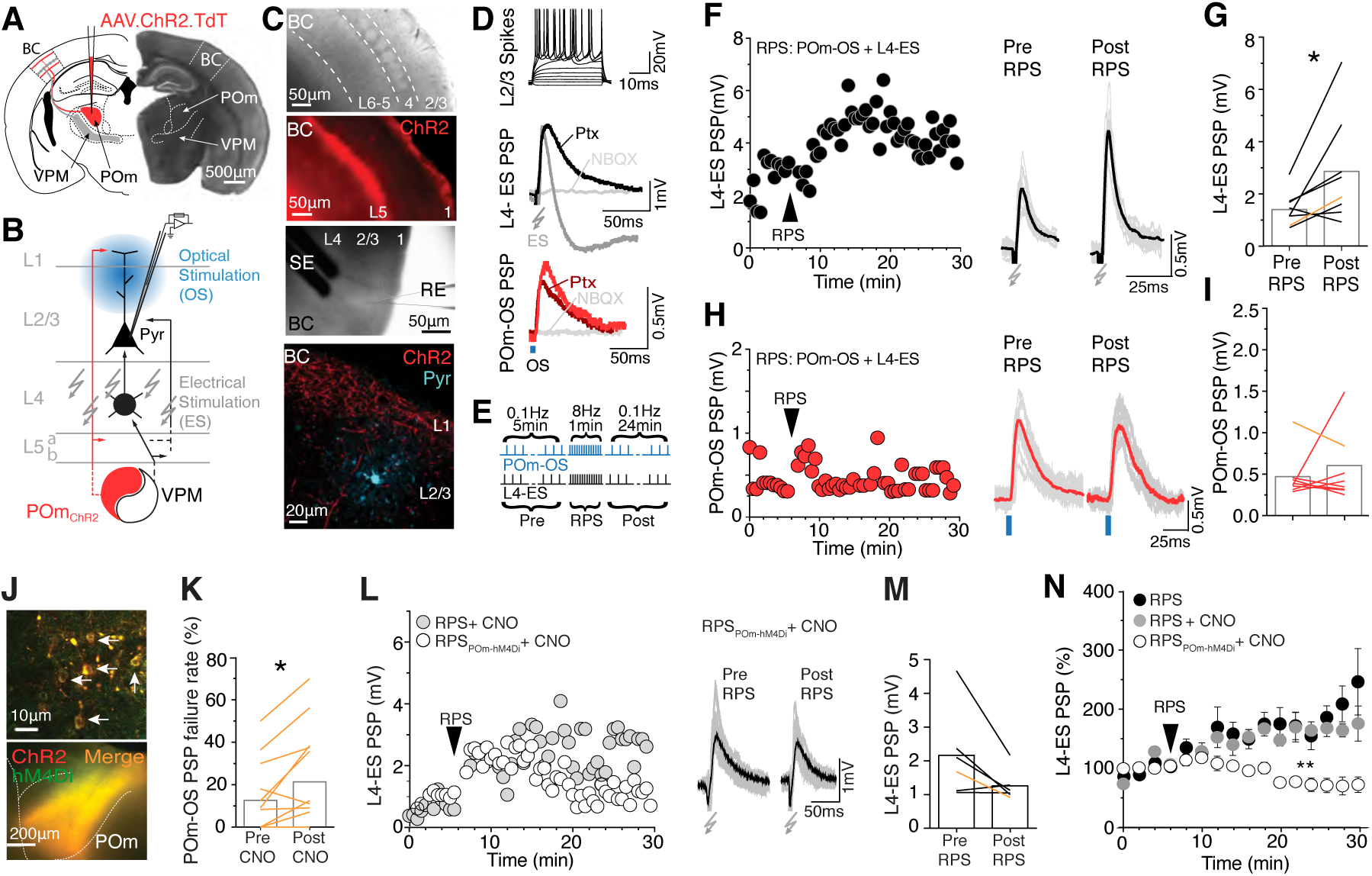
POm inputs facilitate LTP of L2/3 pyramidal neuron synapses. (A) Schematic and bright field image of the POm and VPM and their projections to the barrel cortex (BC) in thalamocortical slices. AAV-mediated expression of ChR2-Tdtomato is directed to the POm. (B) Schematic of the somatosensory thalamocortical (POm and VPM) projections and their relation to intracortical circuits in the BC. Recordings are made in L2/3 pyramidal neurons, while electrically stimulating L4 (L4-ES) and/or optically stimulating the ChR2-expressing POm projections (POm-OS). (C) *Top*, bright field image of the BC. *Middle top*, fluorescent image of L1 and L5 ChR2-tdTomato expressing POm projections. *Middle bottom*, example of experimental configuration; a bipolar stimulating electrode (SE) in L4 and a recording electrode (RE) on a L2/3 pyramidal neuron. *Bottom*, confocal image of ChR2-tdTomato-expressing (ChR2) POm projections in L1 and L2/3, and a L2/3 pyramidal neuron (Pyr) filled with biocytin-streptavidin after patching. (D) *Top*, typical L2/3 pyramidal neuron firing pattern upon current injection steps (40pA). *Below*, Representative traces of L4-ES PSP with/without bath application of Ptx (100μM) and NBQX (10μM). *Bottom*, representative trace of POm-OS PSP with/without Ptx (100μM) and NBQX (10μM). (E) Experimental protocol: alternating L4-ES and POm-OS at 0.1Hz (pre RPS; 5min), followed by rhythmic pairing of L4-ES and POm-OS at 8Hz (RPS; 1min), followed by alternating L4-ES and POm-OS at 0.1Hz (post RPS; 24 min). (F) *Left*, L4-ES PSP amplitudes in an example cell. *Right*, representative L4-ES PSPs pre vs. post RPS. Grey lines represent individual traces, and black lines their average (G) The population (bars) and cell (lines) mean L4-ES PSP amplitudes pre vs. post RPS; n=8 cells; *P*=0.04; paired Student’s t-test. Yellow line, representative cell in (F). (H) *Left*, POm-OS PSP amplitudes in an example cell. *Right*, representative POm-OS PSP pre vs. post RPS. (I) Mean POm-OS PSP amplitudes pre vs. post RPS; n=8 cells; *P*=0.57; paired Student’s t-test. (J) *Bottom*, Confocal image of POm expression of hM4Di-mCitrine (green), ChR2-tdTomato (red). *Top*, magnified confocal image of POm cells expressing both hM4Di-mCitrine and ChR2-tdTomato. (K) POm-OS PSP failure rate (%) pre CNO vs. post CNO; n=13; *P*=0.016; paired Student’s t-test. (L) *Left*, L4-ES PSP amplitudes in example cells for RPS_POm-hM4Di_ and RPS, both with CNO. *Right*, representative L4-ES PSP, pre and post RPS_POm-hM4Di_+CNO. (M) Mean L4-ES PSP amplitudes pre vs. post RPS_POm-hM4Di_+CNO; n=6; *P*=0.07; paired Student’s t-test. (N) Normalized L4-ES PSP amplitudes after RPS under various conditions. RPS+CNO drives LTP, whereas RPS_POm-hM4Di_+CNO fails to elicit LTP; *P*=0.01. The addition of CNO does not alter the ability of RPS to drive LTP; *P*=0.76; two-way repeated measures ANOVA.

In our laboratory it was previously demonstrated that cortical L2/3 pyramidal neurons in S1 undergo post-synaptic LTP following a brief period (1min) of rhythmic whisker stimulation (RWS) (Gambino et al., 2014). This form of LTP does not rely on back-propagating action potentials (bAPs), but is driven by long-lasting N-methyl-D-aspartate receptor (NMDAR)-mediated potentials that are dependent on the activity of the POm. This suggests that lemniscal as well as paralemniscal activity is necessary to induce LTP. However, it remains unclear if co-activity of the POm and L4 alone is sufficient to drive LTP in L2/3 pyramidal neurons, and what, exactly, are the underlying microcircuits within S1 that mediate this LTP.

Here we aimed at dissecting the circuit underpinnings of this type of plasticity in thalamocortical slices by isolating the synaptic inputs that we suspected are driving the RWS-evoked LTP in L2/3 pyramidal neurons *in vivo*. We paired optogenetic stimulation of POm afferents and electrical stimulation of L4 over the same time-course and at the same frequency (1min, 8Hz) as LTP-evoking RWS *in vivo*. We demonstrate that this rhythmic paired stimulation (RPS) of POm-originating and L4-originating pathways can drive LTP of L2/3 pyramidal neuron excitatory synapses. This type of LTP is occluded by prior RWS *in vivo*. Furthermore, we show that the POm provides direct inputs onto VIP interneurons.

The paired stimulation (PS) of L4 and the POm increases their activity, whereas it reduces SST interneuron activity, causing a disinhibiton of L2/3 pyramidal neurons. Finally, we found that both direct POm input to L2/3 pyramidal neurons and the disinhibition are necessary to drive LTP.

Altogether, this study shows a form of LTP in S1 that is mechanistically linked to sensory-driven plasticity. It is dependent on the co-activation of intracortical connections along with higher-order thalamocortical feedback input and is gated by local VIP-mediated disinhibition, revealing a powerful circuit motif for cortical plasticity.

## RESULTS

### Higher-order POm thalamic inputs are indispensible for LTP of intracortical synapses on L2/3 pyramidal neurons

To test if RPS of L4 and the POm can drive synaptic LTP we recorded intracellular responses from L2/3 pyramidal neurons in thalamocortical slices while pairing electrical stimuli (ES) of L4 with optical stimuli (OS) of POm afferents expressing the light-gated ion channel channelrhodopsin-2 (ChR2) at 8Hz for 1 minute (**Figures 1A,B**) (Zhang et al., 2006). ChR2 was expressed in POm neurons using targeted injections of recombinant adeno-associated viral vectors (AAV) encoding ChR2 under the CMV promoter. Successful injections could be identified by virtue of a robust expression of ChR2-tdTomato in POm neurons, as well as by the distinct expression pattern in the barrel cortex of S1, where dense projections could be observed in L1 and L5 and not in L4 (**Figures 1A,C; Supplementary Figure 1**) (Wimmer et al., 2010). Typical spiking patterns induced by current steps identified L2/3 pyramidal neurons (**Figure 1D**) (Avermann et al., 2012).

A single electrical stimulation pulse in L4 (L4-ES, 0.2ms) evoked a depolarizing postsynaptic potential (PSP) in L2/3 pyramidal neurons, incidentally followed by a hyperpolarizing overshoot (**Figure 1D**). The latter component was eliminated by blocking of γ-aminobutyric acid receptors (GABARs) using bath application of picrotoxin (Ptx, 100μM, specifically GABA_A_R). Optical stimulation of ChR2-expressing POm projections (POm-OS, 5ms pulse) consistently evoked a depolarizing PSP (**Figure 1D**). Bath application of Ptx had no effect on the POm-evoked PSP. Bath application of 2,3-dihydroxy-6-nitro-7-sulfamoyl-benzo[f]quinoxaline-2,3-dione (NBQX, 10μM) completely eliminated the L4 and POm-evoked PSPs, indicating dependence on α-amino-3-hydroxy-5-methyl-4-isoxazolepropionic acid receptors (AMPARs, **Figure 1D**).

We performed rhythmic paired stimulation (RPS) of L4 and POm (8Hz, 1min) and measured both L4 and POm-evoked PSP amplitudes pre and post pairing (**Figure 1E**). RPS significantly increased mean L4-evoked PSP amplitudes (**Figure 1F,G**). Mean POm-evoked PSP amplitudes were not significantly potentiated (**Figure 1H,I**), which demonstrates that the LTP is expressed on intracortical and not on thalamocortical synapses.

To determine whether activity of POm afferents is necessary for RPS-driven LTP we repeated the RPS experiment with both ChR2 and hM4Di (inhibitory Designer Drugs Exclusively for Designer Receptors, DREADD) receptors present in the POm (**Figure 1J**) (Armbruster et al., 2007). The hM4Di receptors were activated by bath application of the synthetic agonist clozapine-N-oxide (CNO, 500nM), which diminished the likelihood of eliciting a POm-evoked PSP (59% increase in failure rate; **Figure 1K**) (Stachniak et al., 2014). Under these conditions RPS did not elicit significant LTP (**Figure 1L-M**). This effect was also not attributable to the CNO itself, since the presence of CNO did not prevent RPS-driven LTP in slices that lacked hM4Di expression (**Figure 1L,N**). This suggests that reduced POm activity prevents LTP, which is consistent with previous findings *in vivo* (Gambino et al., 2014). To corroborate these findings we tested the effect of L4 rhythmic electrical stimulation only (L4-RES, 8Hz, 1min; **Supplementary Figure 1**). Mean L4-evoked PSP amplitudes were not significantly increased (**Supplementary Figure 1**). Nonetheless, in 4 out of 7 cells L4-RES induced a significant LTP. These data suggest that L4-RES alone is able to induce LTP in some cells. L4-ES may, however, variably recruit POm ascending fibers passing through L4. Therefore, to eliminate any residual contribution of POm-derived inputs in the L4-RES paradigm, we repeated the experiment using hM4Di expression in the POm. Upon silencing of POm afferents, L4-RES failed to increase the mean L4-evoked PSP amplitudes (**Supplementary Figure 1**). Normalized L4-evoked PSP amplitudes were significantly larger after the L4-RES protocol as compared to L4-RES with POm inhibition (**Supplementary Figure 1**). None of the suppressed LTP effects above were attributable to a change in baseline L4 or POm PSP amplitudes as across experiments baseline PSP size was not correlated with LTP size; nor was there a correlation between LTP size and various electrophysiological parameters (**Supplementary Figure 1**). Together, the data strongly suggests that the activity of POm inputs is required to drive LTP.

We observed no spikes upon RPS or L4-RES. Thus, similar to *in vivo* experiments, the LTP occurs in the absence of bAPs, and instead could have been caused by long-lasting subthreshold depolarization (Gambino et al., 2014). Indeed, we found an increase in cumulative PSP amplitudes upon RPS as compared to L4-RES with the POm inhibited (**Supplementary Figure 1**). The amplitude of the 1^st^ PSP upon the repeated pairing, which is a measure of the increased depolarization was, however, predictive of the size of the LTP (**Supplementary Figure 1**).

Altogether, these data indicate that the activation of POm-derived paralemniscal circuitry is necessary to increase the depolarization of L2/3 pyramidal neurons during the rhythmic stimulation and to potentiate the synapses from intracortical circuits (**Figure 1G,I,M,N Supplementary Figure 1**). Hence, in all of the following experiments we used RPS-driven LTP to investigate the cellular and circuit underpinnings.

### RPS-evoked LTP is NMDA-dependent and shares expression mechanisms with whisker stimulation-evoked LTP *in vivo*

We next used pharmacology to investigate the mechanisms underlying this LTP. Blocking of GABARs with picrotoxin (Ptx, 100μM) induced a robust LTP in all cells (**Figure 2A,B**). L4-evoked PSP amplitudes did not increase without RPS (**Figure 2E**), excluding the possibility that the observed LTP under GABAR block was caused by a ramping up of baseline responses. When the NMDAR blocker (2R)-amino-5-phosphonovaleric acid (APV, 50μM) was added LTP could not be elicited (**Figure 2C-E**). These data indicate that the LTP occurs at excitatory synapses, is NMDAR-dependent, and is not attributable to inhibitory plasticity.

**Figure 2.**
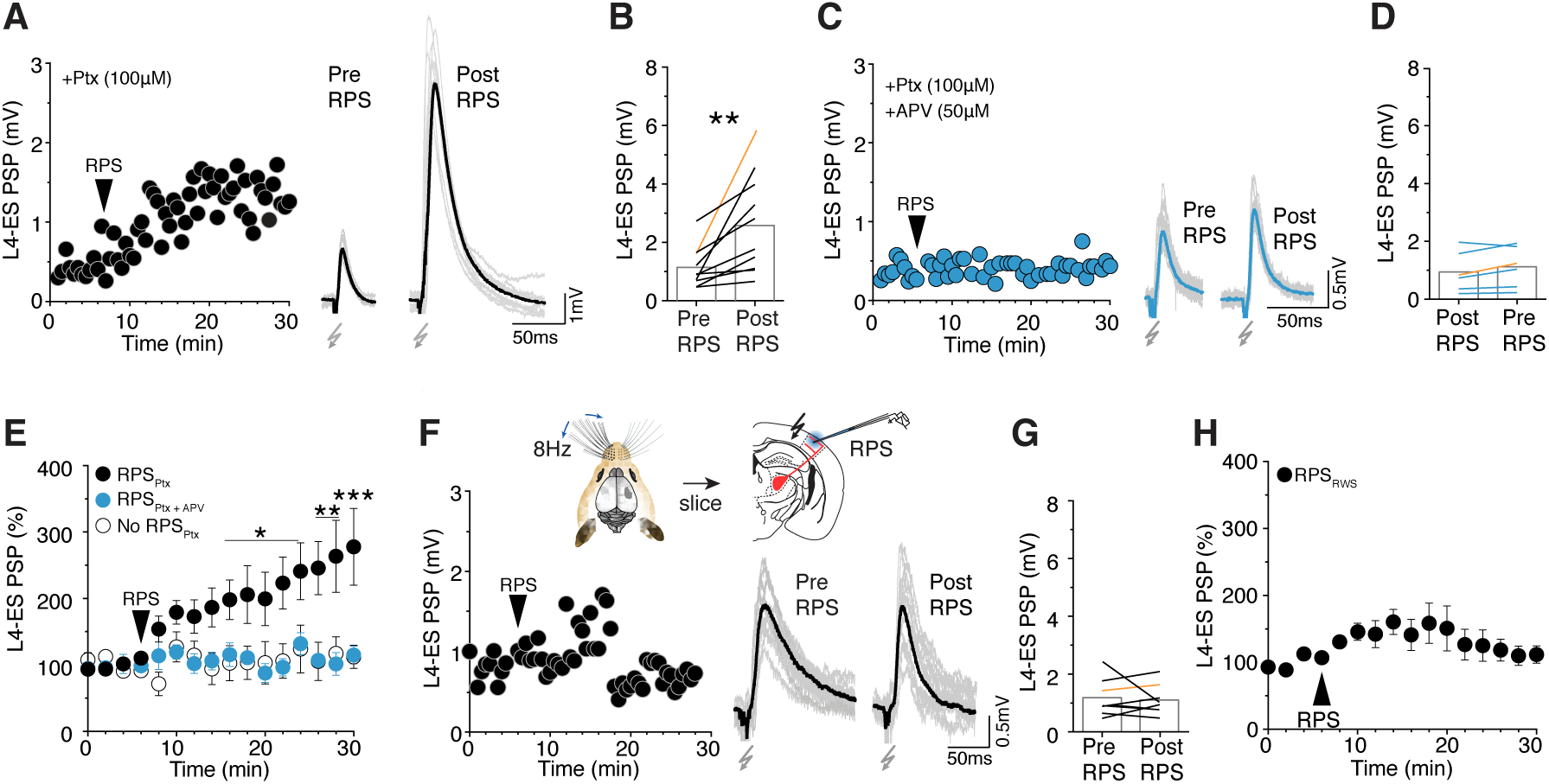
RPS-evoked LTP is NMDA-dependent and shares expression mechanisms with whisker stimulation-evoked LTP *in vivo*. (A) *Left*, L4-ES PSP_Pyr_ amplitudes in an example cell in Ptx (100μM). *Right*, representative L4-ES PSP pre vs. post RPS +Ptx. (B) Mean L4-ES PSP amplitudes pre vs. post RPS +Ptx, n=10 cells, *P*=0.009; paired Student’s t-test. (C) *Left*, L4-ES PSP amplitudes in an example cell +Ptx, APV (50μM). *Right*, Representative L4-ES PSPs pre vs. post RPS +Ptx, APV. (D) Mean L4-ES PSP_Pyr_ amplitudes pre vs. post RPS +Ptx, APV, n=6 cells, *P*=0.11; paired Student’s t-test. (E) Normalized L4-ES PSP amplitudes, comparing RPS + Ptx vs. +Ptx, APV, and No RPS +Ptx (n=5), *P*=0.035; Two-way repeated measures ANOVA. Post-hoc, Bonferroni’s multiple comparisons test, from 22 min: +Ptx vs. +Ptx, APV, *P*<0.02; +Ptx vs. No RPS +Ptx, *P*<0.02; R+Ptx, APV vs. No RPS +Ptx, *P*>0.99). (F) *Top*, Experimental schematic: RWS (all whiskers, 8Hz, 10min) followed by slicing, and RPS (8Hz, 1min). *Below*, L4-ES PSP amplitudes in an example cell for RPS_RWS_. *Right*, representative L4-ES PSP pre and post RPS_RWS_. (G) Mean L4-ES PSP amplitudes pre vs. post RPS_RWS_, n=6 cells, *P*=0.75; paired Student’s t-test. (H) Normalized L4-ES PSP_Pyr_ amplitudes for RPS_RWS._

Similar to the silencing of POm inputs, the NMDAR block reduced PSP amplitudes at the start of the RPS period and significantly impaired the cumulative depolarization (**Supplementary Figure 2**). This is consistent with the *in vivo* observation that POm inputs promote LTP through facilitation of NMDAR-mediated conductances.

We hypothesized that if RPS-driven LTP shares its underlying mechanisms with RWS (rhythmic whisker stimulation)-driven plasticity *in vivo*, RWS would occlude subsequent RPS-driven potentiation in slices from these mice. To test this we rhythmically stimulated all the whiskers with piezoelectric actuators (8 Hz, 10min), a protocol known to induce a robust increase in whisker-evoked cortical local field potentials and LTP (Gambino et al., 2014; Mégevand et al., 2009). This was followed by immediate slice preparation and RPS (RPS_RWS_; **Figure 2F**).

We found that RPS failed to induce LTP in slices of mice that had undergone prior RWS (**Figure 2F-H**). Similarly, RPS-driven LTP in slices followed by a 2^nd^ RPS could not elicit further LTP (**Supplementary Figure 2**).

RWS prior to slicing did not diminish baseline L4-evoked PSP amplitudes, or cumulative PSP amplitudes during RPS_RWS_ (**Supplementary Figure 2**), indicating that the lack of LTP was not due to diminished depolarization, but rather was an effect of occluded expression. This was similarly observed for the repeated RPS slice experiment. Altogether, these results suggest that the paired stimulation of L4 and POm pathways *ex vivo* results in an LTP of the same synapses that are potentiated by RWS *in vivo*, and implies that the same synaptic circuits are recruited by repeated sensory stimuli.

### Paired POm thalamic and L4 cortical inputs engage a disinhibitory microcircuit motif

We next questioned whether the excitatory inputs from L4 and POm onto L2/3 pyramidal neurons are sufficient to induce LTP, or whether local disinhibition is also required. We focused on SST and VIP interneurons. They constitute a well-characterized disinhibitory microcircuit for L2/3 pyramidal cell apical dendrites, which is the location of POm inputs (Wang et al., 2004; Gentet et al., 2012; Pfeffer et al., 2013; Lee et al., 2013). We recorded from these interneurons to determine if they are activated by POm-OS and/or L4-ES, and to measure the effect of paired stimulation (PS) (**Figure 3**).

**Figure 3.**
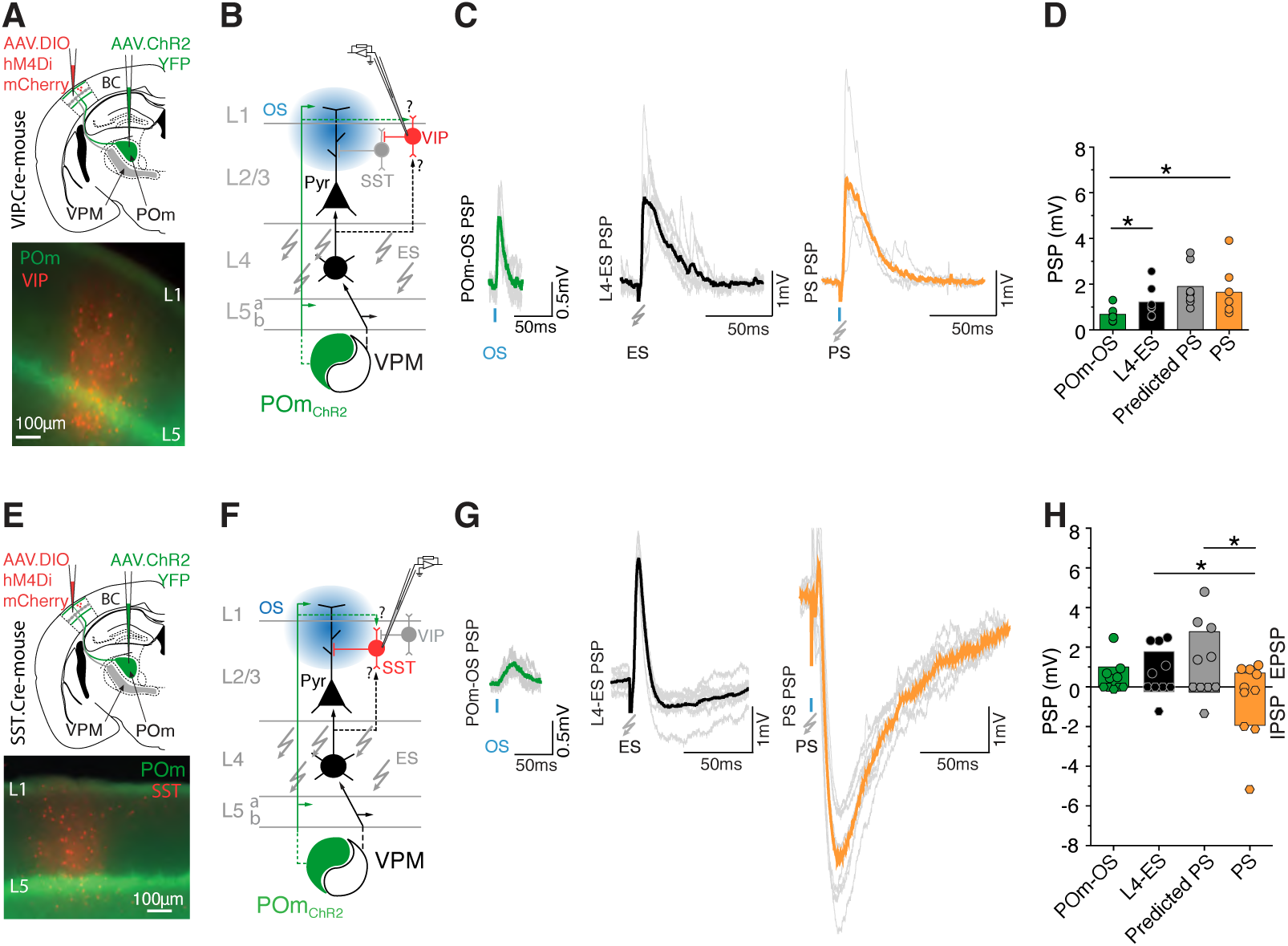
Pairing of L4-ES and POm-OS increases VIP and reduces SST interneuron activity. (A,E) Experimental design and schematic of AAV directed hM4Di-mCherry expression in the BC in a VIP-Cre and SST-Cre driver lines, and ChR2-YFP expression in the POm. *Below*, fluorescence image of hM4Di-mCherry expression and ChR2-YFP-positive POm projections in BC. (B,F) Schematic of the circuit, with a targeted patch recording of a hM4Di-mCherry-postive VIP or SST interneuron to measure possible inputs (dotted arrows) from POm and/or L4. (C,G) Representative traces of L4-ES, POm-OS, and PS in VIP (C) or SST (G) interneurons (without CNO). (D) Mean L4-ES PSP amplitudes in VIP interneurons, n=7 (L4 vs. POm, *P*=0.04; POm vs. PS, *P*=0.04; PS vs. L4, *P*=0.06; PS vs. Predicted PS, *P*=0.17); paired Student’s t-test. (H) Mean L4-ES PSP amplitudes in SST interneurons, n=5 (L4 vs. POm, *P*=0.13; POm vs. PS, *P*=0.37; PS vs. L4, *P*=0.02; PS vs. Predicted PS, *P*=0.01); paired Student’s t-test.

We used VIP-Cre and SST-Cre mice in combination with Cre-dependent AAV viral vectors to target expression of hM4Di-mCherry to VIP and SST interneurons (Taniguchi et al., 2011). In both lines, POm neurons were transfected using AAV-ChR2-YFP viral vectors. Cortical injections of the conditional hM4Di-mCherry vector resulted in robust and widespread expression (**Figure 3A,E**, **Supplementary Figure 3**). To determine efficiency and specificity of labeling we performed immunohistochemistry using anti-SST and anti-VIP antibodies. 100% of the hM4Di-mCherry-postive cells were positive for their respective markers (**Supplementary Figure 3**). Labeled cells were found in all layers, in accordance with described expression patterns (Taniguchi et al., 2011; Pfeffer et al., 2013; Prönneke et al., 2015). Recordings were made from mCherry-expressing cells (without DREADD activation; **Figure 3A,B**) in L2/3. The smaller membrane capacitance (Cm) compared to L2/3 pyramidal neurons further supported that we had targeted interneurons (**Supplementary Figure 3**) (Gertler et al., 2008).

POm and L4-stimulation evoked depolarizing PSPs in both interneuron types (**Figure 3C,D,G,H**). The evoked POm/L4 PSP ratios were larger for VIP interneurons, but not for SST interneurons, as compared to L2/3 pyramidal neurons (**Supplementary Figure 3**). This demonstrates that stimulation of both pathways generates synaptic responses in VIP and SST interneurons, and that POm afferents provide a relatively strong input to VIP interneurons. This result is congruent with the previously observed robust POm-to-VIP monosynaptic responses and weak POm-to-SST polysynaptic responses (Audette et al., 2017).

For VIP interneurons a single paired stimulation (PS, POm-OS and L4-ES) evoked significantly larger mean PSP amplitudes than POm-OS alone and was similar to what would be predicted (predicted PS) based on linear summation of average L4-ES and POm-OS responses alone (**Figure 3C,D**).

In contrast, for SST interneurons PS-evoked depolarizing PSP amplitudes were significantly smaller than the L4-ES and predicted PS amplitudes (**Figure 3G,H**). Mean PS PSP amplitudes were not significantly different from POm-OS. In fact, the response frequently turned into a hyperpolarizing PSP (**Figure 3G,H**). Indeed, the PS/L4-ES EPSP ratios were much smaller in SST interneurons as compared to L2/3 pyramidal neurons and VIP interneurons (**Supplementary Figure 3**). These results suggest that PS inhibits SST interneurons. The diminished depolarization could be due to VIP interneuron-mediated inhibition of SST interneurons, which would translate into diminished SST spiking. This in turn would disinhibit L2/3 pyramidal neurons. Indeed, we found that SST and VIP interneurons intermittently spiked at rest. VIP interneurons tended to increasingly spike upon PS, whereas SST interneurons tended to decrease their spiking activity (**Supplementary Figure 3**).

Altogether, these data show that PS of L4 and POm inputs increases VIP and reduces SST interneuron activity, which is a typical attribute of the VIP-SST-L2/3 disinhibitory microcircuit (Pfeffer et al., 2013; Lee et al., 2013).

### Reduced VIP interneuron activity lowers L2/3 pyramidal neuron PSPs and increases inhibitory conductance

To test whether reduced SST interneurons activity disinhibits L2/3 pyramidal neurons, and whether reduced VIP interneurons activity prevents disinhibiton, we recorded from L2/3 pyramidal neurons while reducing the activity of these hM4Di-expressing interneurons with bath applied CNO (**Figure 4A,E**). We assumed that reducing their activity potentially had widespread effects on barrel column circuits based on the finding that hM4Di-expressing interneurons were found in areas (~750 μm) exceeding the size of barrel-related columns, and because the far majority of each interneuron population was expressing the transgene within the transfected areas (**Supplementary Figure 3**).

**Figure 4.**
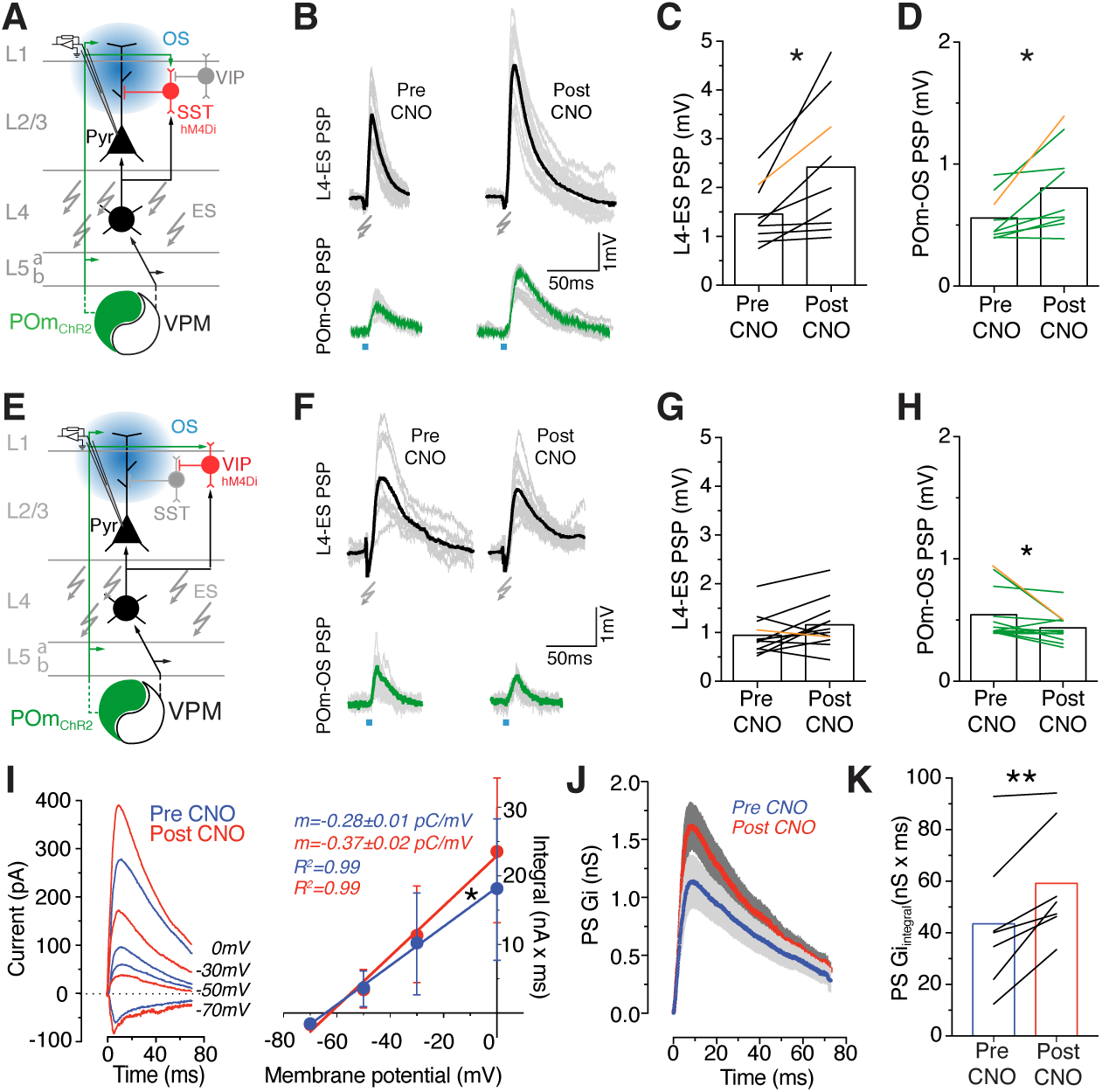
Reduced VIP interneuron activity decreases L2/3 pyramidal neuron PSP amplitudes and increases inhibitory conductances. (A,E) Schematic of the circuit, with a patch recording of L2/3 pyramidal neurons to measure the effects of hM4Di-mediated reduction in SST and VIP interneuron activity. (B,F)*Top*, representative L4-ES PSP pre and post CNO (500nM) in SST-hM4Di (B) and VIP-hM4Di slices (F). *Bottom*, representative POm-OS PSP pre and post CNO in SST-hM4Di (B) and VIP-hM4Di slices (F). (C) Mean L4-ES PSP amplitudes pre vs. post CNO in SST-hM4D slices, n=9 cells, *P*=0.01; paired Student’s t-test. (D) Mean POm-OS PSP amplitudes pre vs. post CNO in SST-hM4Di slices, n=9 cells; *P*=0.02; paired Student’s t-test. (G) Mean L4-ES PSP amplitudes pre vs. post CNO in VIP-hM4Di slices, n=12 cells, *P*=0.06; paired Student’s t-test. (H) Mean POm-OS PSP amplitudes pre vs. post CNO in VIP-hM4Di slices, n=12 cells; *P*=0.04; paired Student’s t-test. (I) *Left*, examples of PS evoked currents in L2/3 pyramidal neurons pre (blue) and post (red) CNO (500nM) in VIP-hM4Di slices at four different holding potentials (−70mV, −50mV, −30mV, and 0mV). *Right*, Synaptic V-I curves (mean±sd). Linearity is assessed by linear regression, slopes pre vs. post CNO, n=7, *P*=0.0365, analysis of covariance (ANCOVA). (J) Averaged PS evoked synaptic inhibitory conductances over time pre vs. post CNO. Shaded areas indicate SEM. (K) mean integrated inhibitory conductance (Gi) in L2/3 pyramidal neurons pre vs. post CNO in VIP-hM4Di slices, n=7 cells, *P*=0.006; paired Student’s t-test.

Firstly, we confirmed that CNO reduced the activity of hM4Di-expressing cells by performing targeted recordings of mCherry-positive neurons. The CNO caused a significant decrease in the resting potential and reduced the ability to induce APs for the same absolute amount of injected current (**Supplementary Figure 4**).

Reduced SST interneuron activity significantly increased L4 and POm-evoked PSP amplitudes in L2/3 pyramidal neurons (**Figure 4B-D**). Conversely, reduced VIP interneuron activity significantly decreased POm-evoked PSP amplitudes (**Figure 4F-H**). This corroborates our hypothesis that the pairing of L4 and POm inputs leads to a disinhibition of L2/3 pyramidal neurons through a POm-to-VIP-to-SST-to-L2/3 microcircuit.

To further confirm that VIP interneurons can disinhibit L2/3 pyramidal neurons upon PS, we performed voltage-clamp recordings in L2/3 pyramidal neurons while silencing VIP interneurons using hM4Di (**Figure 4E**). PS-evoked postsynaptic currents were recorded at various holding potentials (−70mV, −50mV, −30mV, and 0mV) before and after addition of CNO to generate synaptic current-voltage (I-V) curves (**Figure 4I**). Under both conditions we found a linear relationship between the integrated currents and the holding potentials. Reduced VIP interneuron activity significantly increased the slope of the I-V curve (**Figure 4I**). Based on the I-V regression slopes and the synaptic reversal potentials we calculated the inhibitory conductance (Gi) over time (**Figure 4J,K**) (Gambino and Holtmaat, 2012; House et al., 2011; Monier et al., 2008).

The Gi in L2/3 pyramidal neurons significantly increased upon addition of CNO (**Figure 4J,K**). This demonstrates that reduced VIP interneuron activity increases inhibition though other inhibitory interneuron subtypes, most likely though SST interneurons as shown here (Pfeffer et al., 2013), but possibly also though Parvalbumin–expressing (PV) interneurons (Pi et al., 2013). Together these data indicate that increased activity of VIP interneurons as elicited by paired intra-cortical and thalamic POm inputs promotes disinhibition of L2/3 pyramidal neurons. This may gate LTP.

### Reduced VIP interneuron activity prevents RPS-evoked LTP in L2/3 pyramidal neurons

To test whether disinhibiton gates RPS-driven LTP we first measured the effects of RPS on L2/3 pyramidal neurons while reducing the activity of hM4Di-expressing SST interneurons with CNO (**Figure 5A**). Under these conditions RPS evoked significantly larger cumulative PSP amplitudes during rhythmic stimulation as compared to normal RPS (**Supplementary Figure 5**).

**Figure 5.**
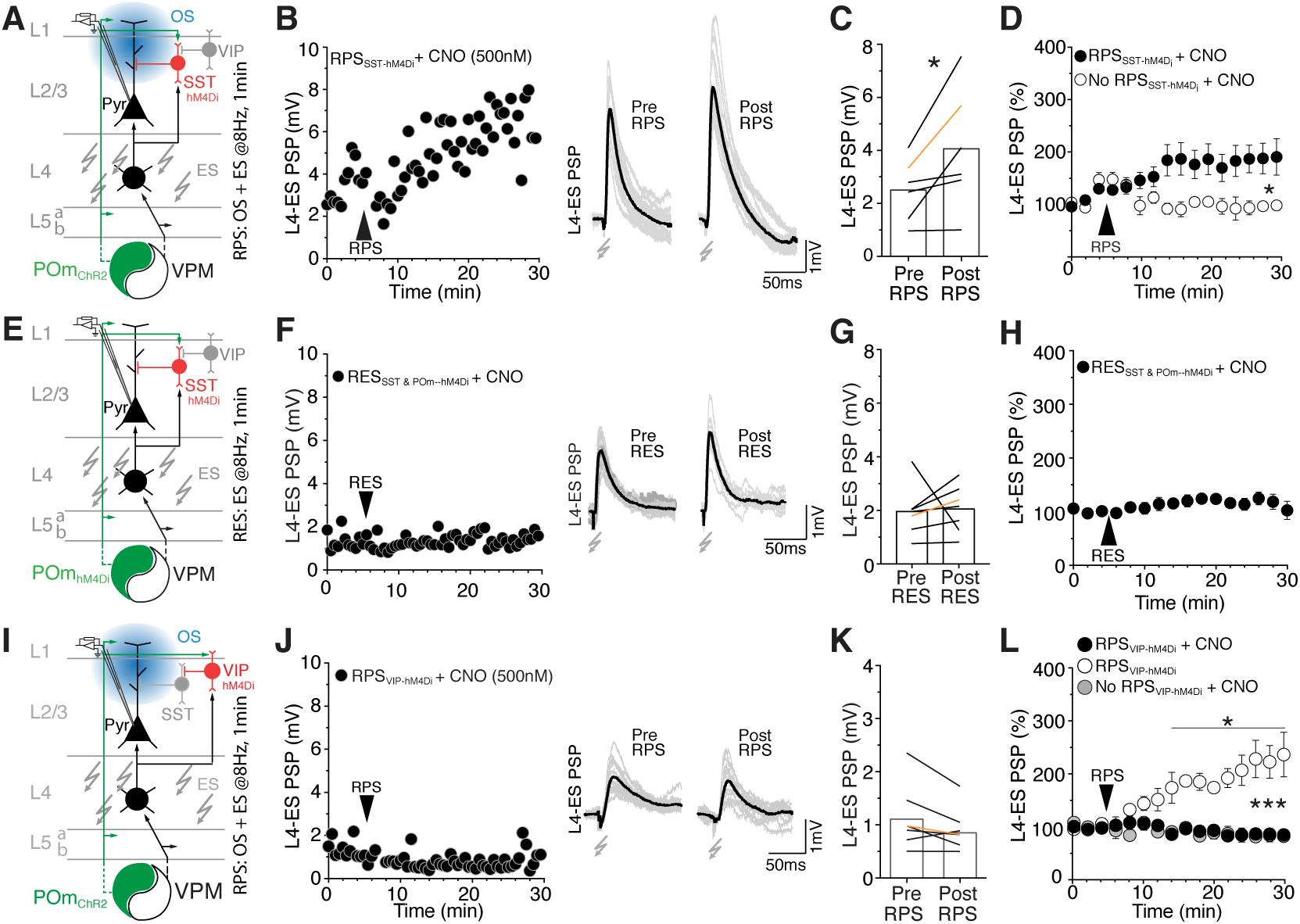
Reduced VIP interneuron activity prevents RPS-evoked LTP in L2/3 pyramidal neurons. (A,E,I) *Top*, schematic of the circuit, with a patch recording of a L2/3 pyramidal neuron to measure the effects of hM4Di-mediated reduction in SST, SST & POm, and VIP activity on RPS or RES-induced plasticity. (B,F,J) *Right*, L4-ES PSP amplitudes in an example cell upon RPS_SST-hM4Di_ (B), RES _SST&POm-hM4Di_ (F), and RPS_VIP-hM4Di_ (J) in CNO. *Left*, representative L4-ES PSPs pre vs. post CNO in RPS_SST-hM4Di_ (B), RES _SST&POm-hM4Di_ (F) and RPS_VIP-hM4Di_ (J) in CNO. (C) Mean L4-ES PSP amplitudes pre vs. post RPS_SST-hM4Di_, n=6 cells, *P*=0.048; paired Student’s t-test. (D) Normalized L4-ES PSP amplitudes, comparing RPS_SST-hM4Di_ to No RPS_SST-hM4Di_ (n=4), *P*=0.049; Two-way repeated measures ANOVA. (G) Mean L4-ES PSP amplitudes pre vs. post RES_SST&POm-hM4Di_, n=7 cells, *P*=0.85; paired Student’s t-test. (H) Normalized L4-ES PSP amplitudes for RES_SST&POm-hM4Di_. (K) Mean L4-ES PSP amplitudes pre vs. post RPS_VIP-hM4Di_, n=7 cells, *P*=0.08; paired Student’s t-test. (L) Normalized L4-ES PSP amplitudes, comparing RPS_VIP-hM4Di_ +CNO, No RPS_VIP-hM4Di_ (n=5 cells), and RPS_VIP-hM4Di_ (n=3 cells), *P*<0.0001 (Post-hoc Bonferroni’s multiple comparisons test, from 12 min: RPS_VIP-hM4Di_ vs. No RPS_VIP-hM4Di_+CNO, *P*>0.99; RPS_VIP-hM4Di_+ CNO vs. RPS_VIP-hM4Di_, *P*<0.03; No RPS_VIP-hM4Di_+CNO vs. RPS_VIP-hM4Di_, *P*<0.007; two-way repeated measures ANOVA).

RPS readily drove LTP under reduced SST interneuron activity (**Figure 5A-C**). Omitting RPS while reducing SST interneuron activity did not increase PSP amplitudes over time, indicating that LTP was not due to a ramping up of responses upon prolonged inactivity (**Figure 5D**). This data is consistent with the idea that disinhibition is a permissive factor for the induction of LTP. This prompts the question as to whether disinhibition alone would be sufficient to drive LTP of rhythmically stimulated intracortical synapses, or whether direct glutamatergic POm input to L2/3 pyramidal neurons is an additional requirement. To test this we expressed hM4Di in SST interneurons as well as in the POm, and reduced both of their activity with CNO while rhythmically stimulating L4 (RES, 8Hz for 1min; **Figure 5E**). RES did not evoke LTP under these conditions. This shows that direct inputs from the POm to L2/3 pyramidal neurons as well as the disinhibiton are required to drive LTP (**Figure 5F-G**).

If VIP interneurons are driving this disinhibiton, and their activation is unequivocally required to facilitate LTP, then reduced VIP interneuron activity should also inhibit the LTP. To test this we measured the effects of RPS on L2/3 pyramidal neurons while reducing the activity of hM4Di-expressing VIP neurons with CNO (**Figure 5I**). Under these conditions RPS resulted in significantly smaller cumulative and mean PSP amplitudes throughout the pairing (**Supplementary Figure 5**), and it did not drive LTP (**Figure 5J-K**). RPS could, however, induce LTP when CNO was not present (**Figure 5L**), and omitting RPS did not increase PSP amplitudes, indicating respectively, that the lack of LTP was not due to the expression of hM4Di per se and not caused by a ramping down of PSP amplitudes with prolonged VIP interneuron inactivation (**Figure 5L**).

Altogether, these data show that the repeated coincident activation of intracortical synaptic circuitry together with higher-order thalamic input gates plasticity of intracortical synapses in S1 via disinhibition.

## DISCUSSION

We showed that the rhythmic co-activation (RPS, 8Hz) of L4-ascending (lemniscal) and POm-feedback (paralemiscal) projections to S1 induces LTP of synapses on L2/3 pyramidal neurons. LTP expression was NMDAR-dependent and not caused by plasticity of inhibitory synapses. It was occluded when, immediately prior to brain slicing, whiskers were stimulated (10min at 8Hz). The latter has been shown previously to induce an LTP of whisker-evoked PSPs (Mégevand et al., 2009; Gambino et al., 2014). This suggests that both LTP paradigms share expression mechanisms and most likely recruit the same synaptic circuits. Thus, the *ex vivo* paradigm that we developed here represents a suitable model for dissecting microcircuits that underlie plasticity of cortical pyramidal neurons *in vivo*.

The LTP was observed at synapses that were recruited by electrical stimulation of L4, but was also critically dependent upon POm activity. L4-RES could drive LTP, but this was less reliable than RPS. Moreover, decreasing POm activity during L4-RES and RPS prevented LTP. This is congruent with findings *in vivo*, where a block of POm activity during RWS prevented LTP expression. Thus, collective recruitment or stimulation of intra-cortical, and long-range axons that ascend through L4, including those originating from the POm, underlies the L4-RES that successfully elicited LTP. These findings imply that this type of plasticity is caused by cooperative synapses, similar to what has been observed in other preparations (Golding et al., 2002; Sjöström and Häusser, 2006; Dudman et al., 2007; Brandalise and Gerber, 2014; Basu et al., 2016).

While the activity of POm projections was required for the increase in L4-evoked PSP amplitudes, their own synapses were not themselves potentiated. This suggests that the strength of POm synapses is saturated or that they lack the molecular mechanisms to express LTP upon this type of paired stimulation (Kotaleski and Blackwell, 2010). Alternatively, the differential effects on POm and L4 inputs may be related to the location of their synapses. The electrical stimulus may recruit various ascending projections traversing through L4, including those originating from L4 and L5 neurons (Feldmeyer, 2012; Petreanu et al., 2009; Lefort et al., 2009). Therefore, the potentiated synapses that are recruited by L4 stimulation may be located, not only on basal dendrites but at various locations along the dendritic tree, including apical dendrites. They may be positioned and perhaps clustered around locations susceptible to compartmentalized calcium events (Kleindienst et al., 2011; Takahashi et al., 2012) whereas POm inputs may not. In addition, local depolarization at these clusters could be amplified by the disinhibitory gate that we have illustrated (Gentet et al., 2012; Pfeffer et al., 2013; Lee et al., 2013; Pi et al., 2013).

The LTP occurred in the absence of somatic spikes since we did not observe any during RPS. Thus, the LTP was dependent upon subthreshold depolarization rather than bAPs, similar to RWS-evoked LTP (Gambino et al., 2014) and hippocampal LTP in slices (Golding et al., 2002; Dudman et al., 2007; Brandalise and Gerber, 2014). Indeed, when we examined the first responses upon RPS we noticed that PSP amplitudes were significantly higher as compared to those evoked by L4 stimulation alone. The size of LTP expression was correlated with the amplitude of these initial RPS-evoked PSPs, but not correlated with the size of baseline PSP amplitudes under various experimental conditions. In addition, an NMDAR block diminished the temporal summation of RPS-evoked dendritic depolarization. Thus, similar to RWS-driven LTP *in vivo*, the potentiation of synapses by RPS was dependent on an NMDAR-dependent sustained increase in postsynaptic depolarization.

In addition to excitatory synaptic inputs to pyramidal neurons, both POm and L4 stimulation evoked PSPs in VIP and SST interneurons, in agreement with recent studies (Wall et al., 2016; Audette et al., 2017). Our experiments did not necessarily distinguish between monosynaptic or polysynaptic inputs. Notably, the direct input of POm axons to SST neurons might be very weak (Wall et al., 2016; Audette et al., 2017). Nonetheless, in our experiments, the activation of both pathways caused spikes in both interneurons, and when the two stimuli were combined, the VIP neurons increased their activity. Interestingly, the SST interneurons experienced a decrease in evoked PSP amplitudes; their spiking rate did not increase and even tended to be lower as compared to L4 stimulation alone. Thus, pairing of the two pathways preferentially activates a cortical circuit that increases VIP and suppresses SST interneuron activity, conceptually similar to responses mediated by whisking in S1 (Lee et al., 2013; Gentet et al., 2012); by reinforcement signals in auditory cortex (Pi et al., 2013); and by locomotion in visual cortex (Fu et al., 2014).

L2/3 pyramidal neuron apical dendrites are strongly inhibited by SST interneurons (Wang et al., 2004; Kapfer et al., 2007), which are in turn inhibited, by VIP interneurons (Pfeffer et al., 2013; Lee et al., 2013). Thus, POm and L4 pairing could reduce the inhibition of L2/3 apical dendrites through the suppression of SST interneuron activity, mediated by increased VIP interneuron activity. Various types of long-range and local inputs have been shown to recruit a similar disinhibitory circuit (Lee et al., 2013; Pi et al., 2013; Fu et al., 2014). In our experiments, the synaptic silencing of SST interneurons increased both POm and L4-evoked PSP amplitudes, and synaptic silencing of VIP interneurons suppressed POm-evoked PSPs. Furthermore, reduced VIP interneuron activity increased inhibitory conductances on L2/3 pyramidal neurons when POm and L4 pathways were paired. Therefore, POm activity not only evokes excitatory responses in L2/3 pyramidal neuron dendrites, but also causes a disinhibition when paired with L4 stimulation.

Our results demonstrate that the recruitment of a VIP interneuron-associated disinhibitory motif is essential for eliciting synaptic plasticity, and strongly suggest that excitatory POm projections provide the necessary input to activate it. The effect of these excitatory long-range projections on plasticity, via their activation of disinhibitory VIP interneurons, bears similarities to the effect of the long-range inhibitory projections from the entorhinal cortex that directly inhibit hippocampal CCK interneurons to enhance plasticity (Basu et al., 2016). This is also similar to disinhibition-mediated plasticity that is caused by increased long-range, cholinergic inputs to the auditory cortex (Letzkus et al., 2011); and the plasticity in the visual cortex caused by running (Fu et al., 2015).

The gating of cortical plasticity by the POm could be widespread. The axonal projections of a single POm neuron to S1 spans large cortical areas (Lu and Lin, 1993; Ohno et al., 2012). Therfore, their activation could unlock a large cortical region for plasticity, thereby allowing receptive field changes beyond a single cortical (barrel) column that are dependent on postsynaptic and NMDA-driven mechanisms (Diamond et al., 1994; Gambino and Holtmaat, 2012).

Higher-order thalamic nuclei such as the POm are thought to provide feedback and contextual information to the primary sensory cortex (Larkum, 2013; Sherman, 2016; Roth et al., 2016). Our data suggest that these feedback signals could gate plasticity in pyramidal neurons and reinforce the synapses of the first-order pathways that convey the principal sensory information. This could be a mechanism for the tuning of cortical synaptic circuits during sensory learning. Interestingly, VIP interneurons in S1 are also activated by projections from the vibrissal primary motor cortex (vM1), which highlights another, now motor related, mechanism for disinhibiting L2/3 pyramidal neurons (Lee et al., 2013). Thus, whisking and contextual sensory feedback could cooperate to powerfully gate synaptic plasticity of L2/3 pyramidal neurons in S1.

## ACKNOWLEDGMENTS

This project was supported by the Swiss National Science Foundation (grant #31003A_153228, #CRSII3_154453), and the International Foundation for Research in Paraplegia. We thank Meaghan Creed, Ronan Chéreau, Stéphane Pages, and Foivos Markopoulos for their technical expertise, and comments on the manuscript and experimental design.

## AUTHOR CONTRIBUTIONS

L.E.W designed and performed the experiments, analyzed the data, and wrote the manuscript. A.H. designed the experiments and wrote the manuscript.

## STAR METHODS

KEY RESOURCES TABLE.

**KEY RESOURCES TABLE**
| REAGENT or RESOURCE | SOURCE | IDENTIFIER |
| --- | --- | --- |
| **Antibodies** |  |  |
| VIP, rabbit polyclonal IgG | Immunostar | #20077, RRID: AB_572270 |
| SST, rat IgG2b, YC7 clone | Merck Millipore. | #MAB354, RRID:AB_2255365 |
| Goat anti-Rat IgG (H+L) Cross-Adsorbed Secondary Antibody, Alexa Fluor 647 | Thermo Fisher Scientific | Cat #:A-21247;, RRID AB_141778. |
| Goat anti-Rabbit IgG (H+L) Superclonal™ Secondary Antibody, Alexa Fluor 647 | Thermo Fisher Scientific | Cat # A32733, RRID AB_2633282 |
| **Bacterial and Virus Strains** |  |  |
| AAV1.CAGGS.Flex.ChR2-tdTomato.WPRE.SV40 | U Penn Vector Core, Pennsylvania | https://www.med.upenn.edu/gtp/vectorcore/ |
| AAV5.EF1a.DIO.hChR2(H134R)-eYFP.WPRE.hGH | Addgene | Addgene20298P |
| AAV9.CMV.PI.Cre.rBG | U Penn Vector Core, Pennsylvania | https://www.med.upenn.edu/gtp/vectorcore/ |
| AAV2.hSyn.HA-hm4D(Gi).IRES.mCitrine | UNC vector core | https://www.med.unc.edu/genetherapy/vectorcore |
| rAAV8.hSyn.DIO.hm4D(Gi).mCherry | Addgene | #44362-AAV2 |
| **Chemicals, Peptides, and Recombinant Proteins** |  |  |
| Clozapine *N*-oxide | Tocris | Cat. No. 4936 |
| Picrotoxin | Tocris | Cat. No. 1128 |
| CNQX | Tocris | Cat. No. 1045 |
| DAP5 | Tocris | Cat. No. 0106/1 |
| Biocytin | Tocris | Cat. No. 3349 |
| QX-314Cl | Tocris | Cat. No. 2313 |
| **Experimental Models: Organisms/Strains** |  |  |
| Mouse/C57BL/6Jrj | Janvier Labs | https://www.janvier-labs.com/rodent-research-models-services/research-models/per-species/inbred-mice/product/c57bl6jrj.html |
| Mouse/SST-IRES-Cre | Jackson Labs | #013044<br>RRID:MGI:4838419 |
| Mouse/VIP-IRES-Cre | Jackson Labs | #010908,<br>RRID:MGI:3054170 |
| **Software and Algorithms** |  |  |
| Ephus software | the Janelia Farm Research Center | http://research.janelia.org/labs/display/ephus: the Janelia Farm Research Center |
| Prism 7 for Mac OS X | GraphPad Software, Inc,<br>La Jolla California, | Version 7.0a, April 2,<br>2016, www.graphpad.com |
| Clampfit 10 | Molecular Devices, LLC | Version 10.8.01<br>http://mdc.custhelp.com |
| Image J | The NIH | https://imagej.nih.gov/ij/ |
| MATLAB | MathWorks | https://www.mathworks.com/ |

### Contact for Reagent and Resource Sharing

Further information and requests for resources and reagents should be directed to and will be fulfilled by the Lead Contact, Anthony Holtmaat

### Experimental Model and Subject Details

#### Animals

All procedures were carried out in accordance with protocols approved by the ethics committee of the University of Geneva and the authorities of the Canton of Geneva. All animals were housed at the University of Geneva’s Animal Care facility under normal light/dark cycles. We used male, C57BL/6J mice and two transgenic Cre-recombinase driver lines, one expressing Cre in SST (*SST-ires-Cre*) interneurons and the other in VIP (*VIP-ires-Cre*) interneurons (Taniguchi et al., 2011). AAV-directed injections were performed at 4 weeks of age and after 2-3 weeks of infection (2.5-3 months of age) mice were euthanized and slice electrophysiology was performed. All transgenes were used as homozygotes.

### Method Details

#### Virus Injection

Mice, aged postnatal days 28-35, were anesthetized using isoflurane (4% with 0.5 1min^−1^ O_2_). Body temperature was maintained at 37°C by a feedback controlled heating pad (FHC). Eye ointment was applied to prevent dehydration and mice were put in a stereotaxic frame. The skin was disinfected with betadine. A burr hole was made in the skull with a pneumatic drill above the region of interest. Injections were targeted to the caudal part of the POm (coordinates from bregma: RC,-2.20mm; ML,-1.20mm: DV,-3.00) and/or the BC (coordinates from bregma: RC, −1.5mm; ML-3.5; Z, −0.4) (Gambino et al., 2014). Expression of ChR2-TdTomato or ChR2-YFP expression was targeted to POm neurons using FLEx AAV vectors (AAV1.CAGGS.Flex.ChR2-tdTomato.WPRE.SV40; AAV5.EF1a.DIO.hChR2(H134R)-eYFP.WPRE.hGH), combined with AAV Cre vectors (AAV9.CMV.PI.Cre.rBG). hM4Di DREADD (Armbruster et al., 2007) was expressed in the POm using non-flex AAVs (AAV2.hSyn.HA-hM4D(Gi).IRES.mCitrine). hM4Di-mCherry was expressed in the VIP-Cre and SST-Cre driver lines using a FLEx AAV vector (rAAV8.hSyn.DIO.hm4D(Gi).mCherry).

The virus was injected (~200nl in the POm and ~100nl in the BC) using a glass pipette attached to a hydraulic manipulator (MMO-220A, Narishigi) at a maximum rate of 100nl min^1^. The solution was allowed to diffuse for at least 10min before the pipette was withdrawn. Once injections were completed the craniotomy was filled with Kwik-Cast (WPI) and the skin re-attached with stainless steel staples (Precise DS15, 3M). In accordance with Swiss Federal laws, analgesia as provided by local lidocaine (1%) application. A subcutaneous injection of buprenorphine (Temgesic,0.05mg kg^−1^) was given to reduce postoperative pain.

#### Thalamocortical slice preparation

2-4 weeks post-viral injections, mice were anesthetized with isoflurane and decapitated. A vibrating microtome was used to prepare 350-μm-thick thalamocortical slices according an acute brain slice method for adult and aging animals (Agmon and Connors, 1991; Ting et al., 2014). Slicing was performed in cold NMDG artificial cerebrospinal fluid solution (aCSF, 300-305 mOSm, pH 7.3), containing the following (in mM): 92 NMDG, 2.5 KCl, 1.25 NaH_2_PO_4_*H_2_O, 30 NaHCO_3_, 20 HEPES, 25 glucose, 2 thiourea, 5 Na-ascorbate, 3 Na-pyruvate, 0.5 CaCl_2_*2H_2_0, 10 MgSO_4_*7H_2_O (Agmon and Connors, 1991; Ting et al., 2014). Slices were then transferred to NMDG aCSF solution at 35°C for 20min, after which slices were immersed in a HEPES aCSF solution (300-305mOSm, pH 7.3), at room temperature, (in mM): 92 NaCl, 2.5 KCl, 1.25 NaH_2_PO_4_*H_2_O, 30 NaHCO_3_, 20 HEPES, 25 glucose, 2 thiourea, 5 Na-ascorbate, 3 Na-pyruvate, 2 CaCl_2_*2H_2_O, and MgSO_4_*7H_2_O. 95% O_2_ + 5% CO_2_ was bubbled though all solutions.

#### Electrophysiology

Whole-cell current-clamp recordings were obtained from patched L2/3 pyramidal neurons or fluorescence-guided targeted patched VIP or SST interneurons. Recordings were performed in freshly prepared aCSF, bubbled with 95% O_2_ + 5% CO_2_, at an osmolarity of 300-305mOsm, containing (in mM): 119 NaCl, 24 NaHCO_3,_ 2.5 KCl, 1.25 NaH_2_PO_4_*H_2_O, 2 MgSO_4_*7H_2_O, 2 CaCl_2_*2H_2_.O, and 12.5 Glucose. Patch pipettes (5-8 mΩ) were filled with 290-295 mOsm internal solution containing (in mM): 135 K-gluconate, 5 KCl, 10 Phosphocreatine, 4 Mg ATP, 0.3 NaGTP, 2.68 (0.1%) Biocytin, and 10mM HEPES. After break-in the cell was allowed to equilibrate for 5 minutes. Membrane capacitance (C_m_) and series resistance (R_s_) was documented. A series of hyperpolarizing and depolarizing step currents (40pA increments) of 500ms duration were applied to measure intrinsic properties and spike patterns of each neuron.

Postsynaptic potentials (PSPs) were evoked by electrical stimulation (0.2 ms) with a bipolar stimulating electrode (matrix tungsten electrode, FHC) placed in L4, and by optical stimulation of ChR2-expressing POm fibers. For optical stimulation, a 5 ms light emitting diode (LED) pulse (excitation λ470nm, Thorlabs, Germany) was applied through the objective above L1. Electrical stimuli were tuned to yield a ~1mV PSP baseline response. The power of the optical stimuli after the objective was kept at ~0.9mW x mm^2^.

In whole-cell current-clamp, the experimental LTP protocol consisted of a 5-min baseline period in which electrical stimuli (L4-ES; 0.1Hz) were alternated with optical stimuli (POm-OS; 0.1Hz) every 1 min, followed by a 1-min period of rhythmic paired stimulation (RPS; 8Hz), and a 30-min plasticity readout period with the same stimuli as during baseline. During RPS L4-ES and POm-OS were applied at the exact same time. We then compared the average amplitude of the PSPs over the baseline (pre RPS, 0-5min; 30 stimulations) with those over the final 5 min of the recordings (pre RPS, 25-30min; 30 stimulations). The level of LTP was calculated per cell, as well as an average over cells. Synaptic responses were monitored before, during, and after the RPS.

Voltage-clamp recordings were made using a cesium-based internal solution: (in mM) 135 cesium methylsulfonate, 4 QX-314Cl, 10 HEPES, 10 Phosphocreatine, 4 Mg-ATP, 0.3 Na-GTP, 3 biocytin, 0.1 spermine, 7.25 pH adjusted with CsOH, 290-295 mOsm).

For chemogenetic silencing experiments, CNO (500 nM) was bath applied 5 min prior to the recordings. CNO remained present during the recordings. CNQX (10μM, Tocris), Ptx (100 μM, Tocris) and/or DAP5 (50μM, Tocris) were applied in a similar way and remained present throughout the recordings.

#### Whisker stimulation

For occlusion experiments, 2-4 weeks post-AAV injection, anesthesia was induced using isofluorane and maintained by IP injection of Medetomidine (Dorbene, 1mg kg^−1^) and Midazolam (Dormicum, 5mg kg −^1^) in sterile NaCl 0.9% (MM-mix). All whiskers were deflected (10min, 8Hz) using a piezolelectric ceramic actuator (PL-series PICMA, Physik Intrumente). A perforated plastic plate was attached to the ceramic plate, through which all whiskers were inserted. The plate remained 4mm away from the skin. The voltage applied to the actuator was set to evoke a whisker displacement of 0.6mm with a ramp of 7-8ms. After whisker stimulation, the mouse was immediately decapitated, which was followed by thalamocortical slice preparation, and RPS.

#### Immunohistochemistry

For immunohistochemical detection and quantification of VIP and SST interneurons, after electrophysiology, mouse thalamocortical brain sections were fixed in 4% PFA (pH 7.4) for 18-24 hours. Slices were then incubated for 1 hour, free floating in a blocking solution of PBS (pH 7.4) containing 0.025% Triton and 5% Bovine Serum Albumin (BSA). After blocking, slices were incubated for 18-24 hours in blocking solution containing primary antibodies (VIP, rabbit polyclonal IgG; SST, rat IgG2b) at a 1:500 dilution (Lee et al., 2013). After incubation in primary antibodies, slices were washed 4 times for 10 minutes in PBS plus 5% BSA at room temperature. They were then incubated for 2 hours in PBS solution containing 5% BSA and the appropriate fluorescence-conjugated secondary antibodies (1:400,). After incubation with secondary antibodies, slices were washed 4 times in PBS at room temperature and placed onto glass slides.

### Quantification and Statistical Analysis

#### ChR2 and hM4Di expression analysis

The VPM and POm are juxtaposed to each other in the thalamus and to control for any spill over of virus into the VPM a post-hoc analysis of the BC was performed. Fluorescent images (10x objective) were taken of slices, PFA-fixed immediately after the ephys recordings. Fluorescence was observed in L1 (from the pia 0-200μm staining) and L5 (600-800μm) in all experimental slices. An intensity measurement was performed across the BC. Slices with an intensity measurement of more then 3 x 10^4^ a.u. in L4 (400-600μm) were deemed to have spill-over of AAV in the VPM, and were eliminated from any further LTP analysis.

To estimate the extent of hM4Di-expression, (**Supplementary Figure 3**) visibly positive cells were counted, and expressed as the total number in 100μm increments from the pia, as well as the number within a layer (**Supplementary Figure 3**), as described^48^. Layers were determined from their distance from the pia.

#### Confocal microscopy and immunohistochemical analysis

Images were generated using a confocal laser-scanning fluorescence microscope at 40x magnification fluorescence intensity was measured by delineating the edges of all visible cells using ImageJ software and by calculating mean fluorescence in these regions of interest (ROI).

To avoid counting false-positives, two controls were performed. First, images were taken in an area adjacent to injection area (i.e., cells that were not visibly expressing hM4Di-mCherry; **Supplementary Figure 3**). ROIs were drawn around anti-SST or anti-VIP positive cells, and fluorescence intensity in the red channel was quantified (green data points in **Supplementary Figure 3**). Next, images were taken of the injection area in sections on which only the secondary antibody Alexa 647 was applied. ROIs were drawn around hM4Di-mCherry-positive cells, and fluorescence intensity in the green channel was quantified (red data points in **Supplementary Figure 3**). Each of these quantifications yielded a mean fluorescence ± 2SD, which was subsequently used as the lower-limit on which we based the overlap estimate (i.e. #true positives/#total). Intensities of the experimental cells (yellow data points in **Supplementary Figure 3**) below these limits were considered as false positive in either channel.

#### Data analysis

All relevant raw data are available from the authors. Electrophysiological data were acquired using a Multiclamp 700B Amplifier (Molecular Devices) using Matlab-based Ephus software. The data were Bessel-filtered during the recording at 10 kHz. Offline analysis was performed using Event Detection/Template Matching tools in Clampfit 10 software. Templates were created by extracting and averaging segments of data that were manually identified as corresponding to an event within 5ms of ES and/or OS. The same template was used for all depolarizing PSPs and another was adopted for hyperpolarizing PSPs. In Event Detection/Template Matching, the template is slid along the data trace one point at a time and scaled and offset to optimally fit the data at each point. Optimization of the fit was found by minimizing the sum of the squared errors between the fitted template and the data. Since background noise rarely exceeded four times the standard deviation of the noise this was used for optimum template matching. If the event detection program found an event within the corresponding window following stimulation, the event was manually accepted and the program would calculate the peak amplitude. Data points were removed on the bases of a significant change in Rs throughout the experiment and if post-hoc viral transfection was not specific. Randomization and blinding methods were not used. Data is presented throughout as mean ±SEM unless otherwise stated.

Paired stimulation synaptic conductances were determined using published methods in voltage clamp using PS postsynaptic currents (PSCs) recorded at 4 different holding potentials (−70, −50, −30, and 0mV; 5 PSCs per V; 0.1 Hz) (House et al., 2011; Monier et al., 2008; Gambino and Holtmaat, 2012). The relationship between the synaptic current (*Isyn*) and synaptic conductance (*Gsyn*) ware given by the following equation:

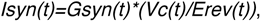

Where *Erev* and *Vc* are the synaptic reversal and holding potential, respectively. For each time point, *Gsyn* and *Erev* are provided by the slope and the x-intercept of the linear regression fit of the I-V curve, respectively. The inhibitory (Gi) conductance was calculated using the following equation:

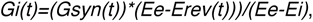

Where *Ee* and *Ei* are the excitatory and inhibitory reversal potential respectively. They were estimated to be −84mV and 0mV, respectively, based on the Nernst Equation, with a 32°C bath temperature, and the internal and external patch solution ion concentrations.

#### Statistical Analysis

For all experiments, n equals the number of cells (no more than 3 cells per mouse per experiment). For immunohistochemical experiments, 3 slices were used from each mouse. All statistical analysis was performed and graphs were created using Prism 7. Unless stated otherwise, a Student’s t-test was used for statistical comparisons. For analysis of data with unequal variances (as determined by a post-hoc F-test), a Mann-Whitney U test was used. For analysis of pre versus post comparisons a paired Student’s t-test was performed. For comparisons over time a Two-way repeated, analysis of variance (ANOVA) was utilized followed by a post-hoc, Bonferroni’s multiple comparisons test. For comparisons of the V-I curves linear regressions were performed and an Analysis of Covariance (ANCOVA) to compare slopes (Figure 4I). Results were considered statistically significant when the P-value < 0.05. No statistical methods were used to estimate sample size. β-power values were calculated and are provided in the Supplementary Information.

---

**Supplementary Figure 1.**
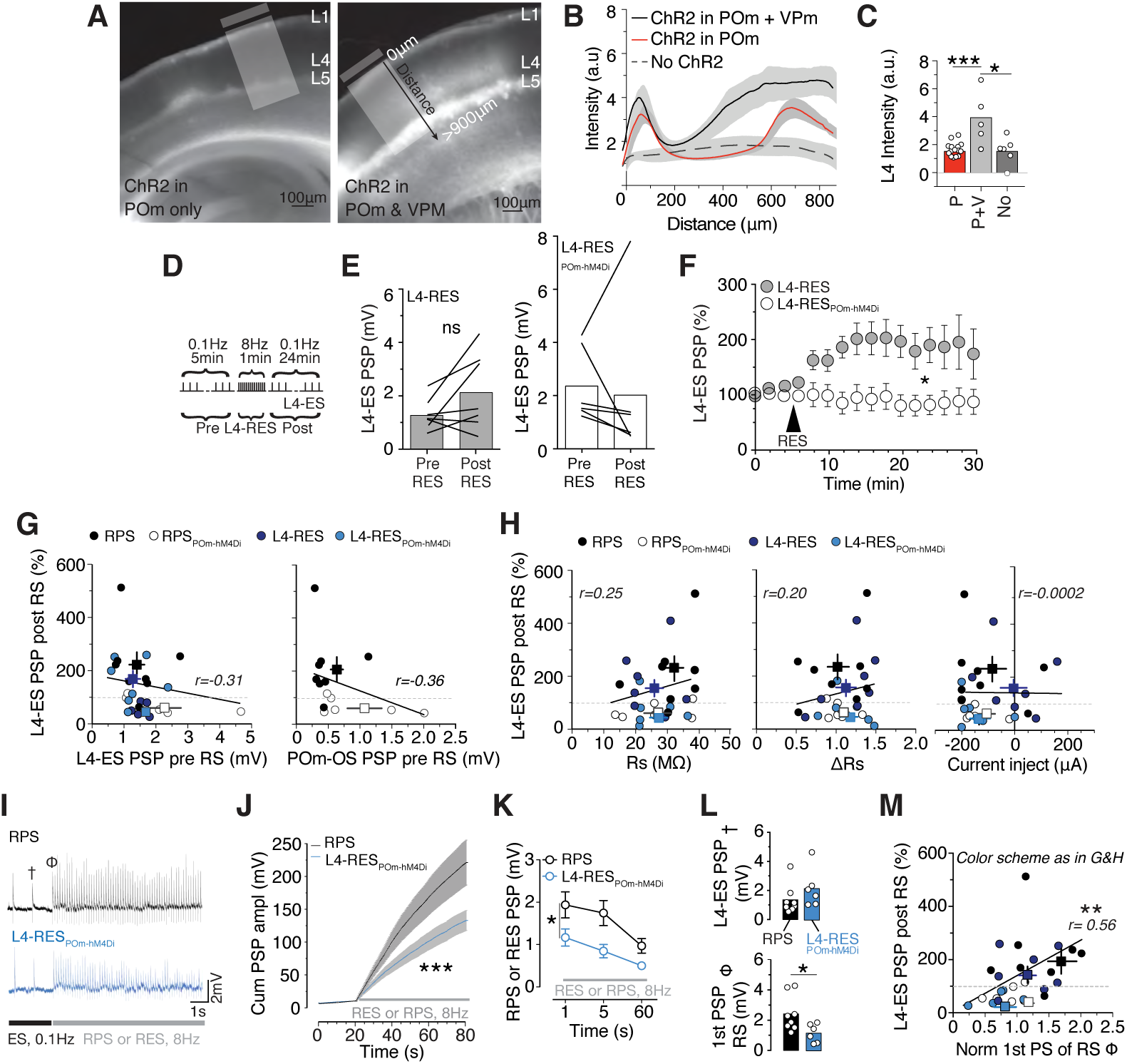
Analysis of AAV-directed expression of ChR 2 and inhibition of the POm prevents RPS induced LTP. (A) Images of thalamocortical slices containing only ChR2-tdTomato-positive POm projections (left) or containing ChR2-tdTomato-positive POm and VPM projections (right) in the BC. (B) Intensity profile (a.u.) of ChR2-tdTomato expression from the pia to L5, comparing POm only, POm + VPM, and No ChR2 expression. (C) Mean L4 (400-600 μM from pia edge) ChR2-tdTomato fluorescence intensity in POm only (P; n=15, 1.5±0.1a.u.), POm + VPM (P+V; n=5, 3.9±0.8a.u.) and No ChR2 (No; n=6, 1.6±0.4a.u). Stats: P vs. P+V, *P*=0.0005 (β=0.99), No vs. P, *P*=0.47. No vs. P+V, *P*=0.018 (β=0.10), Student’s t-tests. (D) *Left*, L4-RES experimental protocol. (E) *Left*, mean L4-ES PSP amplitudes pre (1.27±0.23mV) vs. post (2.23±0.55mV) L4-RES. Stats: n=7, *P*=0.12 (β=0.32), paired Student’s t-test. *Right*, mean L4-ES PSP amplitudes pre (2.36±0.56mV) vs. post (2.02±1.17mV) L4-RES_POm-hM4Di_ + CNO (500nM), n=6, *P*=0.72; paired Student’s t-test (β=0.06). (F) Normalized L4-ES PSP amplitudes, 2 min bins, comparing L4-RES, n=7, vs. L4-RES_POm-hM4Di_, n=6; *P*=0.028; Two-way repeated measures ANOVA. (G) *Left*, mean L4-ES PSP amplitude before rhythmic stimulation (pre RS) vs. LTP size (post RS). Stats: Pearson’s r=-0.31, r^2^=0.096, *P*=0.12. *Right*, mean POm-OS PSP amplitude before RS vs. LTP size. Stats: Pearson’s r=-0.36, r^2^=0.13, *P*=0.20. (H) Rs (*left*), ΔRs (*middl*e), and maximum current injection (*right*) vs. LTP size. Stats: respectively, Pearson’s r=0.25, r^2^=0.6, *P*=0.21; r=0.20, r^2^=0.04, *P*=0.33; r=-0.01, r^2^<0.01, *P*=0.99). (I) Representative traces for the initial portion of the rhythmic (8Hz) stimulation, comparing RPS to L4-RES_POm-hM4Di_. (J) Cumulative PSP amplitudes during rhythmic stimulation. Stats: n=14, *P*<0.0001, two-way repeated measures ANOVA. (K) PSP amplitudes across time points during rhythmic stimulation, comparing RPS to L4-RES_POm-hM4Di_. Stats: n=14, *P*=0.036; two-way repeated measures ANOVA. (L) *Top*, mean L4-ES PSP amplitude at baseline, comparing RPS (1.35±0.36mV, n=8) to L4-RES_POm-hM4Di_ (2.17±0.54mV, n=6). Stats: *P*=0.22 (β=0.97), Student’s t-test. *Bottom*, amplitude of the 1^st^ PSP during the rhythmic stimulation (RS), comparing RPS (2.41±0.42mV, n=8) to L4-RES_POm-hM4Di_ (1.17±0.25mV, n=6). Stats: *P*=0.037 (β=0.99), Student’s t-test. (M) 1^st^ PSP amplitude of RS, normalized to the mean baseline L4-ES PSP amplitude vs. LTP size (%). Stats: Pearson’s r=0.56, r^2^=0.31, *P*=0.003.

**Supplementary Figure 2.**
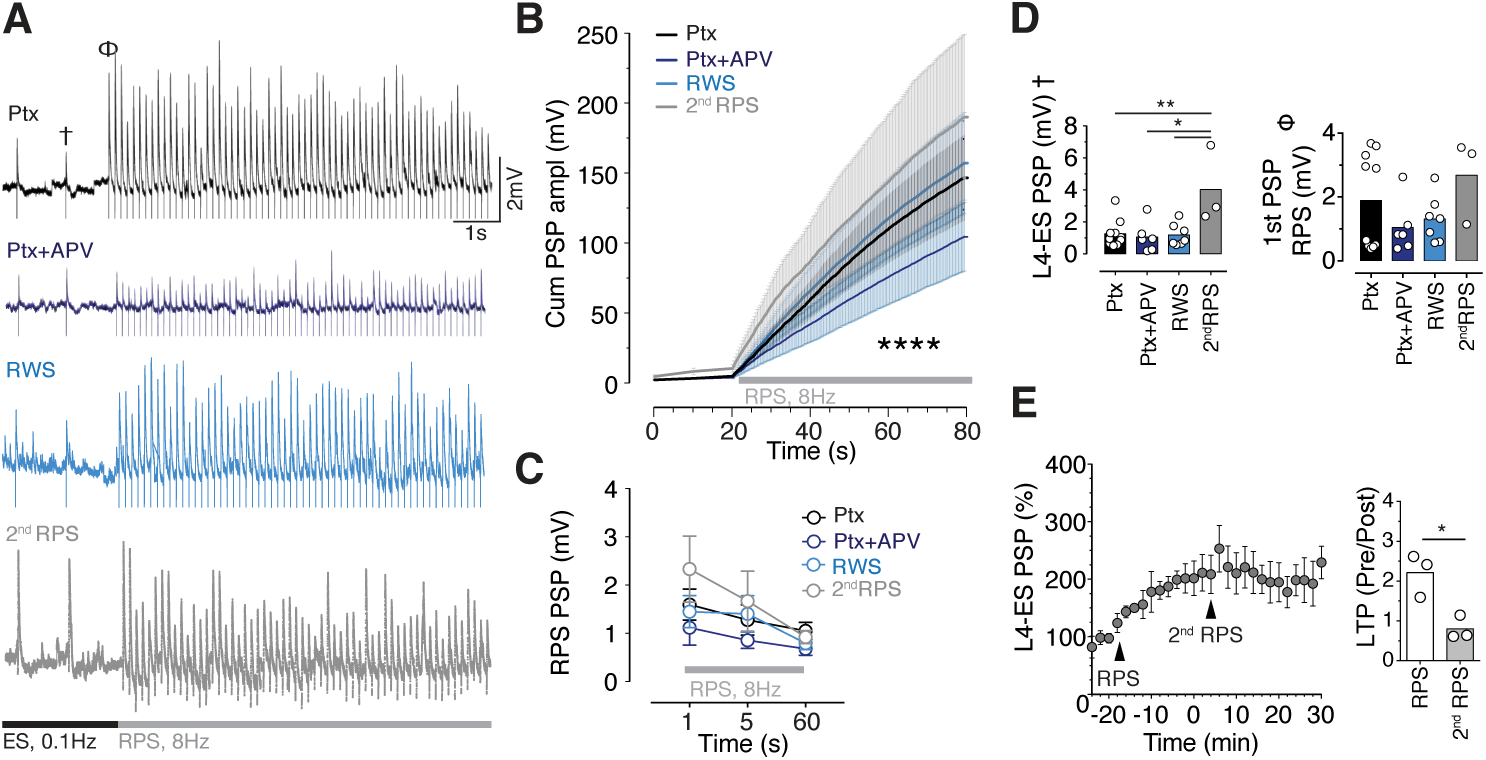
RPS-induced cumulative PSP amplitudes under various conditions. (A) Representative traces for the initial portion of rhythmic (8Hz) stimulation, comparing recordings under Ptx, Ptx+APV, RWS, and a 2^nd^ RPS. (B) Cumulative PSP amplitudes during RPS, comparing Ptx, Ptx+APV, RWS, and a 2^nd^ RPS. Stats: *P*<0.0001, Two-way ANOVA. Post-hoc Bonferroni’s multiple comparisons: Ptx vs. Ptx+APV, *P*<0.0001; Ptx vs. RWS, *P*=0.04; Ptx vs. 2^nd^ RPS, *P*<0.0001; Ptx+APV vs. RWS, *P*<0.0001; RWS vs. 2^nd^ RPS, *P*<0.0001. (C) Mean PSP amplitude across time points during RPS, comparing Ptx, Ptx+APV, RWS, and 2^nd^ RPS. Stats: *P*=0.49; repeated measures Two-way ANOVA. (D) *Left*, Mean L4-ES PSP amplitude at baseline, comparing Ptx (n=10, 1.25±0.27mV), Ptx+APV (n=6, 1.05±0.38mV), RWS (n=7, 1.18±0.26mV), and 2^nd^ RPS (n=3, 4.02±1.39mV). Stats: Ptx vs. Ptx+APV, *P*=0.67 (β=0.23); Ptx vs. RWS, *P*=0.87 (β=0.25); Ptx+APV vs. RWS, *P*=0.77 (β=0.25); 2^nd^ RPS vs. Ptx, *P*=0.008 (β=0.99); 2^nd^ RPS vs. Ptx, *P*=0.03 (β=0.98); 2^nd^ RPS vs. RWS, *P*=0.02; Student’s t-tests. *Right*, 1^st^ PSP amplitudes during RPS, comparing Ptx (n=10, 1.89±0.47mV), Ptx+APV (n=6, 1.04±0.33mV), RWS (n=7, 1.31±0.27mV), and 2^nd^ RPS (n=3, 2.69±0.77mV). Stats: Ptx vs. Ptx+APV, *P*=0.23 (β=0.98); Ptx vs. RWS, *P*=0.36 (β=0.90); Ptx+APV vs. RWS, *P*=0.55 (β=0.45); 2ndRPS vs. Ptx, *P*=0.43 (β=0.45); 2^nd^ RPS vs. Ptx+APV, *P*=0.05; 2^nd^ RPS vs. RWS, *P*=0.06 (β=0.45); Student’s t-tests. (E) Normalized L4-ES PSP amplitudes across RPS followed by a 2ndRPS. (G) LTP ratio (pre/post) comparing the 1^st^ RPS (n=3, 2.21±0.31) to the 2ndRPS (0.80±0.17). Stats: *P*=0.02 (β=0.99); Student’s t-test.

**Supplementary Figure 3.**
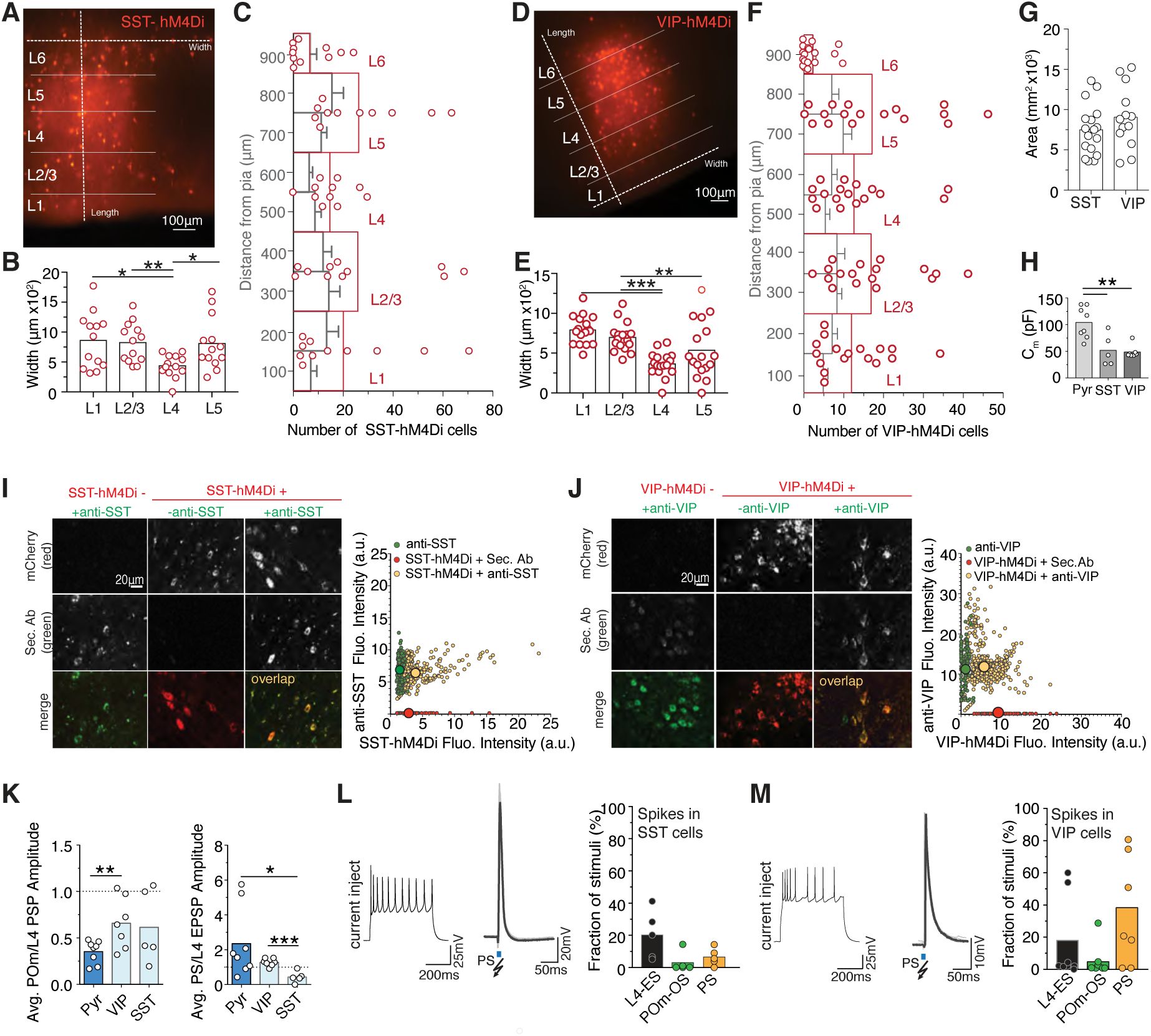
Analysis of h M 4 Di-mCherry expression and spiking in SST and VIP-Cre driver lines. (A,D) Image of SST-hM4Di (A) and VIP-hM4Di (D) expression in L1-L6 of the BC. Dotted lines indicate the dimensions over which the area of expression was measured. (B) Width measurement of expression across layers in the SST-Cre driver line. Stats: L1 (n=13, 859±462μm) vs. L4 (436±191μm), *P*=0.04 (β=1.00); L2/3 (824±331μm) vs. L4, *P*=0.001(β=0.19); L4 vs. L5 (809±440μm), *P*=0.01(β=1.0); and L2/3 vs. L5, *P*=0.94; Student’s t-tests. (C) Number of cells per layer expressing hM4Di-mCherry in the SST-Cre driver line vs. distance from the pia (100μm binning). (E) Width measurement of expression across layers in the VIP-Cre driver line. Stats: L1 (n=17, 792±180μm) vs. L4 (364±152μm), *P*<0.0001; L2/3 (697±178μm) vs. L4, *P*<0.0001 (β=1.0); L4 vs. L5 (536±355μm), *P*=0.20 (β=1.0); L1 vs. L5, *P*=0.015 (β=1.0); and L2/3 vs. L5, *P*=0.06 (β=0.99); and L1 vs. L2/3, *P*=0.13 (β=0.99); Student’s t-tests. (F) Number of cells per layer expressing hM4Di-mCherry in the VIP-Cre driver line vs. distance from the pia (100μm binning). (G) hM4Di expression area in SST-Cre mice (n=14, 0.90±0.10mm^2^×10^3^) compared to VIP-Cre mice (n=15, 0.72±0.08mm^2^×10^3^). Stats: *P*=0.17 (β=1.0); Student’s t-test. (J) Membrane capacitance (C_m_), comparing Pyr (n=15, 99±8pF), SST (n=5, 52±13pF) and VIP (n=7, 49±4pF) cells. Stats: Pyr vs. SST, *P*=0.008 (β=1.0); Pyr vs. VIP, *P*=0.001 (β=1.0); VIP vs. SST, *P*= 0.79 (β=0.13); Student’s t-tests. (I) *Left*, confocal images of anti-SST and SST-hM4Di-mCherry overlap in the injection area (SST-hM4Di+ & anti-SST+), outside of the injection area (SST-hM4Di-& anti-SST+), and in control sections with secondary antibody only (SST-hM4Di+ & anti-SST-). *Right*, fluorescence intensity in the injection areas (yellow dots) SST-hMD4i Fluo. Intensity (3.7±0.1a.u.) vs. anti-SST Fluo. Intensity (6.4±0.1a.u.), outside the injection area (green) SST-hMD4i (1.6±0.3a.u) vs. anti-SST (6.9±0.1a.u), and in control slices with secondary antibody only (red) SST-hMD4i (2.9±0.2a.u.) vs. anti-SST (0.2±0.01a.u). In the SST-line (n=7), 77% of the anti-SST-positive cells co-expressed hM4Di-mCherry (i.e. efficiency), the other 33% were anti-SST positive cells but not transfected. 100% of the hM4Di-mCherry-postive cells were labeled with anti-SST (i.e. specificity). (J) *Left*, confocal images of anti-VIP and VIP-hM4Di-mCherry overlap in the injection areas (VIP-hM4Di+ & anti-VIP+), outside of the injection area (VIP-hM4Di-& anti-VIP+), and in control sections with secondary antibody only (VIP-hM4Di+ & anti-VIP-). *Right*, Fluorescence intensity in the injection area (yellow) VIP-hMD4i Fluo. Intensity (5.9±0.2a.u.) vs. anti-VIP Fluo. Intensity (12.6±0.2a.u.), outside of the injection area (green) VIP-hMD4i (0.9±0.01a.u.) vs. anti-VIP (12.7±0.5a.u.), and in control sections with secondary antibody only (red) VIP-hMD4i (9.5±0.2a.u.) vs. anti-VIP (0.2±0.04 a.u.). In the VIP-line (n=8), 86% of the anti-VIP-positive were found to co-express hM4Di-mCherry, and 100% of the hM4Di-mCherry-postive cells were labeled with anti-VIP. (K) *Left*, POm-OS over L4-ES PSP amplitude ratios, comparing Pyr (n=8, 0.35±0.04), VIP (n=7, 0.66±0.1), and SST (n=5, 0.62±0.2). Stats: Pyr vs. VIP *P*=0.001 (β=1.0); Pyr vs. SST, *P*=0.4 (β=0.99); VIP vs. SST, *P*=0.83 (β=0.99) paired Student’s t-tests. *Right*, PS over L4-ES PSP amplitude ratios, comparing Pyr (n=8, 2.35±0.7), VIP (n=7, 1.23±0.08), and SST (n=5, 0.43±0.4). Stats: Pyr vs. VIP *P*=0.40 (β=1.0); Pyr vs. SST, *P*=0.01 (β=1.0); VIP vs. SST, *P*=0.0003 (β=0.99); paired Student’s t-tests. (L,M) *Left*, typical spiking pattern of a SST (L) and VIP (M) interneurons after a depolarizing current step. *Middle*, representative trace of a spike upon PS. *Right*, Fraction of stimuli (%) that induced spikes in SST (L) and VIP (M) interneurons. Stats: comparing spikes in SST cells: POm (n=5, 3.1±2.8%) vs PS (6.6±2.4%), *P*=0.5 (β=0.78); POm vs. L4 (20.13±6.76), *P*=0.1(β=0.99); PS vs. L4, *P*=0.07 (β=0.99); comparing spikes in VIP cells, POm (n=7, 7.1±11.8%) vs. PS (34.2±13.4%), *P*=0.07 (β=0.99); POm vs. L4 (17.77±10.16%), *P*=0.35 (β=0.72); PS vs. L4, *P*=0.038 (β=0.93); paired Student’s t-tests.

**Supplementary Figure 4.**
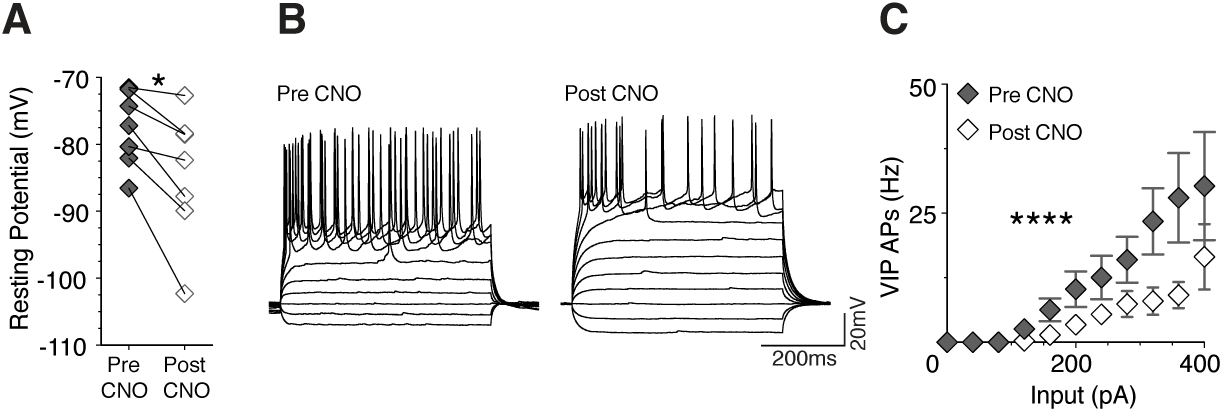
Validation of hM4Di DREADDs in VIP interneurons. (A) Resting membrane potential of VIP interneurons. Stats: pre CNO (n=7, −77.6±2.12mV) vs. post CNO (84.56±3.70mV), *P*=0.01 (β=0.99), paired Student’s t-test. (B) Representative traces of hyperpolarizing and depolarizing current steps (40pA) pre and post CNO in VIP interneurons. (C) AP frequency (Hz) as a function of current input (pA). Stats: n=7, *P*<0.0001, two-way ANOVA.

**Supplementary Figure 5.**
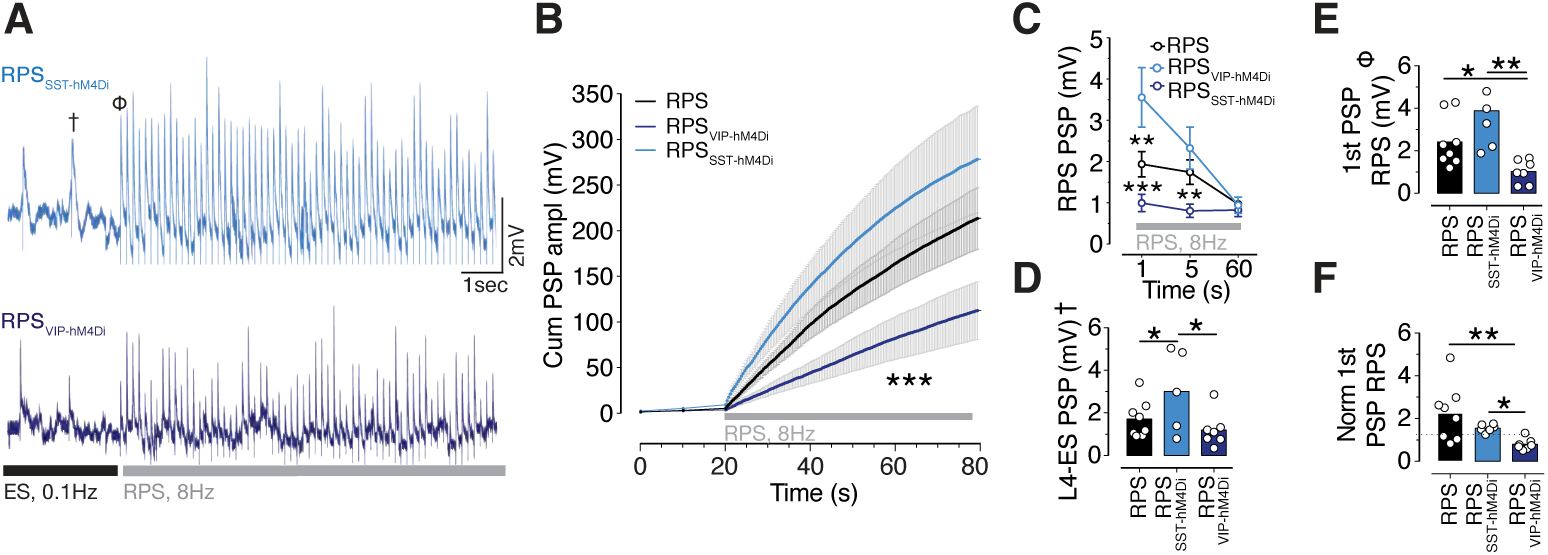
SST and VIP interneurons bi-directionally modulate RPS-induced cumulative PSP amplitudes in L2/3 pyramidal neurons. (A) Representative traces for the initial portion of RPS while reducing SST (RPS_SST-hM4Di_) or VIP (RPS_VIP-hM4Di_) interneuron activity. (B) Cumulative PSP amplitudes in L2/3 pyramidal neurons during RPS, comparing RPS, RPS_SST-hM4Di_ and RPS_VIP-hM4Di_. Stats: n=21, *P*<0.0001; two-way ANOVA. Post-hoc Bonferroni’s multiple comparisons: RPS vs. RPS_VIP-hM4Di_, *P*<0.0001; RPS vs. RPS_SST-hM4Di_, *P*<0.0001; RPS_VIP-hM4Di_ vs. RPS_SST-hM4Di_, *P*<0.0001. (C) Mean PSP amplitude across time points during RPS, comparing RPS, RPS_VIP-hM4Di_, and RPS_SST-hM4Di_. Stats: *P*=0.01, two-way repeated measures ANOVA. Post-hoc Bonferroni’s multiple comparisons: 1sec, RPS vs. RPS_VIP-hM4Di_, *P*=0.12; RPS vs. RPS_SST-hM4Di_, *P*=0.003; RPS_VIP-hM4Di_ vs. RPS_SST-hM4Di_, *P*<0.0001. 5sec, RPS vs. RPS_VIP-hM4Di_, *P*=0.12; RPS vs. RPS_SST-hM4Di_, *P*=0.65; RPS_VIP-hM4Di_ vs. RPS_SST-hM4Di_, *P*>0.99. 60sec, RPS vs. RPS_VIP-hM4Di_, *P*>0.99; RPS vs. RPS_SST-hM4Di_, *P*=0.65; RPS_VIP-hM4Di_ vs. RPS_SST-hM4Di_, *P*>0.99. (D) Mean L4-ES PSP amplitudes at baseline, comparing RPS (n=8, 1.41±0.24 mV), RPS_VIP-hM4Di_ (n=7, 1.13±0.23mV), and RPS_SST-hM4Di_ (n=6, 2.51±0.48mV). Stats: RPS_VIP-hM4Di_ vs. RPS_SST-hM4Di_, *P*=0.02 (β=0.99); RPS_VIP-hM4Di_ vs. RPS, *P*=0.42 (β=0.99); RPS_SST-hM4Di_ vs. RPS, *P*=0.045; Student’s t-tests. (E) 1^st^ RPS PSP amplitude, comparing RPS (2.41±0.42mV), RPS_VIP-hM4Di_ (1.02±0.21mV), and RPS_SST-hM4Di_ (3.89±0.79mV). Stats: RPS vs. RPS_VIP-hM4Di_ *P*=0.01 (β=1.00); RPS vs. RPS_SST-hM4Di_, *P*=0.12 (β=0.99); RPS_VIP-hM4Di_ vs. RPS_SST-hM4Di_, *P*=0.02; Student’s t-tests. (F) 1^st^ RPS PSP amplitude, normalized to mean baseline L4-ES PSPs, comparing RPS (2.41±0.42mV), RPS_VIP-hM4Di_ (1.02±0.21 mV), and RPS_SST-hM4Di_ (3.89±0.79mV). Stats: RPS vs. RPS_VIP-hM4Di_ *P*=0.004 (β=1.00); RPS vs. RPS_SST-hM4Di_, *P*=0.12 (β=0.99); RPS_VIP-hM4Di_ vs. RPS_SST-hM4Di_, *P*=0.02 (β=1.00); Student’s t-tests.

## Supplementary Values and Statistics

***Figure 1G:*** Mean L4-ES PSP_Pyr_ amplitude pre (1.41±0.24mV) vs. post (2.87±0.72mV) RPS; n=8 cells; *P*=0.0426; paired Student’s t-test (β= 0.57).

***Figure 1I :*** Mean POm-OS PSP_Pyr_ amplitude before pre (0.47±0.1mV) vs. post (0.56±0.15 mV) RPS; n=8 cells; *P*=0.566; paired Student’s t-test (β=0.082).

***Figure 1J:*** Mean L4-ES PSP_Pyr_ amplitude pre (2.17±0.54mV) vs. post (1.26±0.22mV) RPS; n=6 cells; *P*=0.0709; paired Student’s t-test (β=0.06).

***Figure 1K:*** POm-OS PSP failure rate (%) pre CNO (12.64±4.58%) vs. post CNO (21.36±6.56%); n=13; *P*=0.016; paired Student’s t-test (β=0.72).

***Figure 1M:*** Mean L4-ES PSP amplitudes, pre (2.17±0.54mV) vs. post (1.26±0.19mV) RPS_POm-hM4Di_ + CNO; n=6; *P*=0.07; paired Student’s t-test (β=0.46).

***Figure 1N:*** Normalized L4-ES PSP amplitudes after RPS under various conditions. RPS+CNO vs. RPS_POm-hM4Di_+CNO fails to elicit LT; *P*=0.01. RPS vs. RPS+CNO *P*=0.76; two-way repeated measures ANOVA.

***Figure 2B:*** Mean L4-ES PSP_Pyr_ amplitude pre (1.14±0.64mV) vs. post (2.87±0.72mV) RPS_Ptx_; n=10 cells; *P*=0.0088; paired Student’s t-test (β=0.93).

***Figure 2D:*** Mean L4-ES PSP_Pyr_ amplitude pre (0.95±0.70 mV) vs. post (1.12± 0.70mV) RPS_Ptx, APV_; n=6 cells; *P*=0.113; paired Student’s t-test (β=0.35).

***Figure 2E:*** Normalized L4-ES PSP amplitude (%), 2 min bins, RPS_Ptx_ vs. RPS_Ptx, APV_ vs. No RPS_Ptx_ n=5 cells, *P*=0.0351; Two-way repeated measures ANOVA. Post-hoc, Bonferroni’s multiple comparisons test, from 22mins: RPS_Ptx_ vs. RPS_Ptx, APV_ *P*>0.021; RPS_Ptx_ vs. No RPS_Ptx_, *P*>0.02; RPS_Ptx, APV_ vs. No RPS_Ptx_, *P*>0.9999.

***Figure 2G:*** Mean L4-ES PSP amplitude RPS_RWS_ pre (0.95±0.70mV) vs. post (1.12±0.70mV); n=7 cells; *P*=0.7503; paired Student’s t-test (β=0.42).

***Figure 2H:*** Normalized L4-ES PSP amplitude (%), 2min bins, for RPS_RWS._

***Figure 3D:*** Mean PSP amplitudes in VIP interneurons comparing L4 (1.23±0.27mV) vs. POm (0.68±0.12mV), n=7 cells, *P*=0.04 (β=0.58); POm (0.68±0.12mV) vs. PS (1.65±0.43mV), n=7 cells, *P*=0.04 (β=0.59); PS vs. L4, n=7 cells, *P*=0.06 (β =0.60); POm vs. Predicted PS (1.9±0.36mV), *P*=0.004 (β=0.96); L4 vs. Predicted PS, P=0.001(β=0.99); PS vs. Predicted PS, n=7 cells, *P*=0.17 (β=0.26); paired Student’s t-tests.

***Figure 3H:*** Mean PSP_SST_ amplitudes comparing L4 (1.78±0.37mV) vs. PS (0.70±0.19mV), n=5 cells, *P*=0.013 (β=0.83); PS vs. Predicted PS (2.78±0.63mV), n=5 cells, *P*=0.01 (β=0.88); PS vs. POm (1.00±0.38mV), n=5 cells, P=0.37 (β=0.26); Predicted PS vs. L4, n=5 cells, *P*=0.0563 (β=0.52); Predicted PS vs. POm, n=5 cells, *P*=0.0089 (β=0.52); paired Student’s t-tests.

***Figure 4C:*** Mean L4-ES PSP_Pyr_ amplitude pre CNO (1.46±0.20mV) vs. post (2.42±0.46 mV); n=9 cells; *P*=0.0135; paired Student’s t-test (β=0.79).

***Figure 4D:*** Mean POm-OS PSP_Pyr_ amplitude pre CNO (0.56±0.063mV) vs. post (0.80±0.135); n=9 cells; *P*=0.0224; paired Student’s t-test (β=0.70).

***Figure 4G:*** Mean L4-ES PSP_Pyr_ amplitude pre CNO (0.94±0.12mV) vs. post (1.16±0.14mV); n=12 cells; *P*=0.0639; paired Student’s t-test (β=0.47).

***Figure 4H:*** Mean POm-OS PSP_Pyr_ amplitude pre CNO (0.54±0.06mV) vs. post (0.44±0.04mV); n=9 cells; *P*=0.0445; paired Student’s t-test (β=0.54).

***Figure 4I :*** Slopes pre (−0.28±0.01pC/mV) vs. post CNO (−0.37±0.02pC/mV), n=7, *P*=0.0365, an Analysis of Covariance (ANCOVA).

***Figure 4K:*** Mean pre CNO (43.53±10.12nS) vs. post CNO (59.14±8.466nS), n=7 cells, *P*=0.0064, paired Student’s t-test (β=0.92)

***Figure 5C:*** Mean L4-ES PSP amplitude pre RPS_SST-hM4Di_ (2.51±0.48mV) vs. post (4.06±0.94mV); n=6 cells, *P*=0.0478; paired Student’s t-test (0.56).

***Figure 5d:*** Normalized L4-ES PSP_Pyr_ peak amplitude (%), 2 min bins, comparing RPS_SST-hM4Di_ + CNO, n=6 cells, vs. No RPS_SST-hM4Di_ + CNO, n=4 cells, *P*=0.0493; Two-way repeated measures ANOVA.

***Figure 5G:*** Mean L4-ES PSP amplitudes pre (1.96±0.36mV) vs. post (2.053±0.33mV). RES_SST&POm-hM4Di_, n=7 cells, *P*=0.8477; paired Student’s t-test (β=0.05).

***Figure 5K:*** Mean L4-ES PSP amplitude pre RPS_VIP-hM4Di_ (1.13±0.23mV) vs. post (0.92±0.15mV), n=7 cells; *P*=0.0824; paired Student’s t-test (β=0.42).

***Figure 5L:*** Normalized L4-ES PSP amplitude (%), 2 min bins, comparing RPS_VIP-hM4Di_+ CNO vs. No RPS_VIP-hM4Di_ (n=5 cells) + CNO vs. RPS_VIP-hM4Di_ (n=3 cells)*, P*<0.0001, Two-Way repeated measures ANOVA. Post-hoc Bonferroni’s multiple comparisons test, from 12 min: RPS_VIP-hM4Di_+ CNO vs. No RPS_VIP-hM4Di_, *P*>0.9999; RPS_VIP-hM4Di_+ CNO vs. RPS_VIP-hM4Di_, *P*<0.0295; No RPS_VIP-hM4Di_ vs. RPS_VIP-hM4Di_, *P*<0.0073.

## HIGHLIGHTS

- Higher-order (HO) thalamocortical inputs aid intracortical synaptic plasticity
- HO thalamic inputs increase VIP and decrease SST interneuron activity
- The activation of VIP interneurons disinhibits L2/3 pyramidal neurons
- This novel HO-to-VIP disinhibitory motif gates intracortical synaptic plasticity

## eTOC Blurb

- Using *ex vivo* patch-clamp recordings, optogenetics, and chemogenetics Williams and Holtmaat dissect the circuits underlying sensory-driven LTP in cortex *in vivo*. This reveals a novel circuit motif in which higher-order thalamocortical input gates plasticity of intracortical synapses via VIP-mediated disinhibition.

## CELL PRESS DECLARATION OF INTERESTS POLICY

Transparency is essential for a reader’s trust in the scientific process and for the credibility of published articles. At Cell Press, we feel that disclosure of competing interests is a critical aspect of transparency. Therefore, we ask that all authors disclose any financial or other interests related to the submitted work that (1) could affect or have the perception of affecting the author’s objectivity, or (2) could influence or have the perception of influencing the content of the article, in a “Declaration of Interests” section.

### What types of articles does this apply to?

We ask that you disclose competing interests for all submitted content, including research articles as well as front matter (e.g., Reviews, Previews, etc.) by completing and submitting the “Declaration of Interests” form below. We also ask that you include a “Declaration of Interests” section in the text of all research articles even if there are no interests declared. For front matter, we ask you to include a “Declaration of Interests” section only when you have information to declare.

### What should I disclose?

We ask that you and all authors disclose any personal financial interests (examples include stocks or shares in companies with interests related to the submitted work or consulting fees from companies that could have interests related to the work), professional affiliations, advisory positions, board memberships, or patent holdings that are related to the subject matter of the contribution. As a guideline, you need to declare an interest for (1) any affiliation associated with a payment or financial benefit exceeding $10,000 p.a. or 5% ownership of a company or (2) research funding by a company with related interests. You do not need to disclose diversified mutual funds, 401ks, or investment trusts.

### Where do I declare competing interests?

Competing interests should be disclosed on the “Declaration of Interests” form as well as in the last section of the manuscript before the “References” section, under the heading “Declaration of Interests”. This section should include financial or other competing interests as well as affiliations that are not included in the author list.

Examples of “Declaration of Interests” language include:

> “AUTHOR is an employee and shareholder of COMPANY.”
>
> “AUTHOR is a founder of COMPANY and a member of its scientific advisory board.”

*NOTE:* Primary affiliations should be included on the title page of the manuscript with the author list and do not need to be included in the “Declaration of Interests” section. Funding sources should be included in the “Acknowledgments” section and also do not need to be included in the “Declaration of Interests” section. (A small number of front-matter article types do not include an “Acknowledgments” section. For these articles, reporting of funding sources is not required.)

### What if there are no competing interests to declare?

For *research* articles, if you have no competing interests to declare, please note that in a “Declaration of Interests” section with the following wording:

> “The authors declare no competing interests.”

*Front-matter* articles do not need to include this section when there are no competing interests to declare.

## CELL PRESS DECLARATION OF INTERESTS FORM

If submitting materials via Editorial Manager, please complete this form and upload with your final submission. Otherwise, please e-mail as an attachment to the editor handling your manuscript.

Please complete each section of the form and insert any necessary “Declaration of Interest” statement in the text box at the end of the form. A matching statement should be included in a “Declaration of Interest” section in the manuscript.

### Institutional Affiliations

We ask that you list the current institutional affiliations of all authors, including academic, corporate, and industrial, on the title page of the manuscript. ***Please select one of the following:***

- All affiliations are listed on the title page of the manuscript.
- I or other authors have additional affiliations that we have noted in the “Declaration of Interests” section of the manuscript and on this form below.

### Advisory/Management and Consulting Positions

We ask that authors disclose any position, be it a member of a Board or Advisory Committee or a paid consultant, that they have been involved with that is related to this study. ***Please select one of the following:***

- The authors have no positions to declare.
- I or other authors have management/advisory or consulting relationships noted in the “Declaration of Interests” section of the manuscript and on this form below.

### Patents

We ask that you disclose any patents related to this work by any of the authors or their institutions. ***Please select one of the following:***

- The authors have no related patents to declare.
- I or one of my authors have a patent related to this work, which is noted in the “Declaration of Interests” section of the manuscript and on this form below. Please include patent number(s).

***Please insert any “Declaration of Interests” statement in this space*.** This exact text should also be included in the “Declaration of Interests” section of the manuscript. If no authors have a competing interest, please insert the text, “The authors declare no competing interests.”

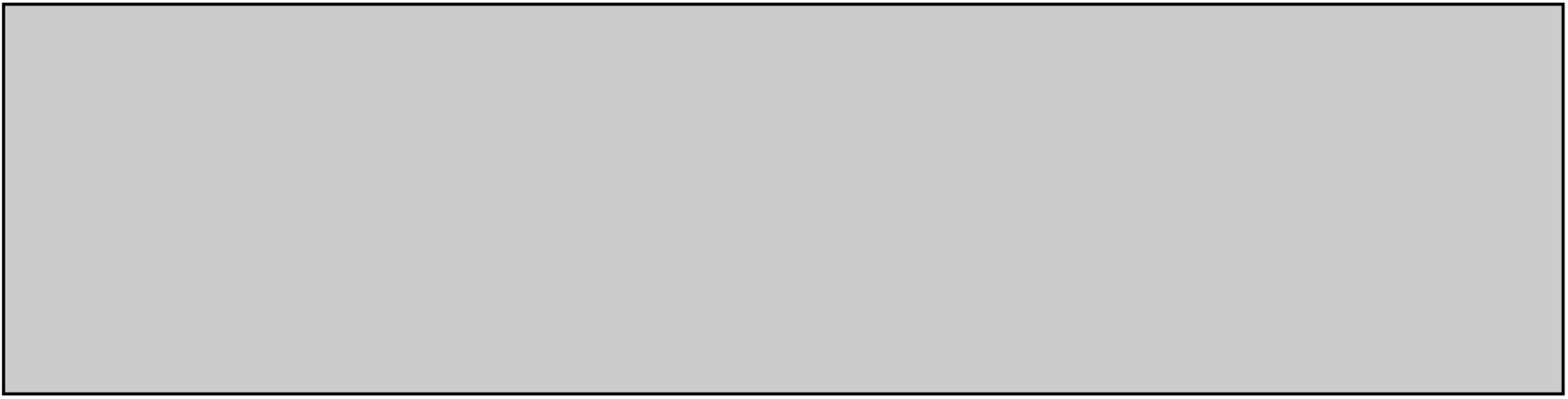

**On behalf of all authors, I declare that I have disclosed all competing interests related to this work. If any exist, they have been included in the “Declaration of Interests” section of the manuscript.**

**Name:** Anthony Holtmaat

**Manuscript Number (if available):**

